# Initial tumor composition shapes resistance evolution and treatment outcomes in non-small cell lung cancer

**DOI:** 10.64898/2026.08.31.748215

**Authors:** Mohammadreza Satouri, Joel S. Brown, Jafar Rezaei, Kateřina Staňková, Rachel Cavill

## Abstract

Drug resistance is a leading cause of treatment failure in non-small cell lung cancer (NSCLC), yet how resistance evolves during treatment and whether its fitness consequences depend on tumor composition remains poorly understood. Using a game-theoretic mathematical model fitted to longitudinal *in-vitro* data from alectinib-sensitive and alectinib-resistant H3122 NSCLC cells grown under different treatment and microenvironmental conditions, we found that the fitness effect of evolving resistance depended critically on the initial proportion of resistant cells in the tumor. When resistant cells were initially rare, resistance evolved faster and increasing resistance was associated with a growth advantage. When resistant cells were initially frequent, increasing resistance was associated with a fitness cost. In both cases, increasing resistance eroded treatment efficacy. In the gain-of-resistance regime, stabilization therapy could maintain a stable tumor equilibrium only if resistant cells were excluded. Maximum tolerated dosing was not always optimal for maximizing time to progression; intermediate doses performed better when they kept the initial tumor growth rate close to zero. These results suggest that evolutionary therapy for NSCLC should account not only for the abundance of resistant cells, but also for how resistance is evolving and what fitness consequences it currently carries in individual patients.

**Significance:** Evolving resistance in NSCLC can either increase or decrease resistant-cell fitness depending on initial tumor composition. Evolutionary therapy should therefore account for resistance evolution, not only resistant-cell abundance.

## 1 Introduction

Therapy resistance is a leading cause of treatment failure in advanced cancer, and is particularly relevant in non-small cell lung cancer (NSCLC), where initially effective targeted therapies are frequently followed by recurrent or progressive disease [1–5]. Intratumor heterogeneity further complicates treatment because cancer cell subpopulations can differ in proliferation rate, interactions among each other, and sensitivity to therapy [6–8]. Evolutionary therapy aims to exploit this heterogeneity, and to forestall or delay resistance by anticipating and steering cancer dynamics [4, 9, 10]. A common principle underlying such strategies is that treatment-sensitive cells can suppress resistant cells through competition when therapy is reduced or temporarily withdrawn [11–13]. This principle is especially useful when resistance carries a fitness cost in the absence of therapy: resistant cells survive treatment better, but grow more slowly than sensitive cells when treatment pressure is low [12, 14]. Under this assumption, evolutionary therapy can use sensitive cells as a competitive resource to delay resistant-cell expansion and prolong tumor control.

However, resistance need not have a fixed effect on cellular fitness. Resistance mechanisms may impose metabolic or physiological costs, for example through drug-efflux activity, altered DNA repair, or changes in target pathways [15–17]. Yet resistant cells may also compensate for these costs, acquire additional growth advantages, or benefit from conditions created by treatment [18–20]. Resistance can also be dynamic rather than fixed: during treatment, resistant cells may continue to change genetically, epigenetically, or phenotypically, altering both their growth rate and their sensitivity to therapy [21–23]. This raises a clinically important question for evolutionary therapy: how does evolving resistance affect resistant-cell fitness, treatment efficacy, and the feasibility of tumor control?

Mathematical models provide a way to formalize this question and to address it using longitudinal data. Many evolutionary therapy models distinguish sensitive and resistant populations but treat resistance as a fixed cellular state with a constant effect on fitness and treatment response [11, 12, 24–26]. Recent mathematical approaches have begun to relax these assumptions by allowing resistance traits and treatment response to vary [22, 27–31]. These approaches are particularly relevant for NSCLC, where experiments have shown that treatment and microenvironment can strongly alter the interaction between sensitive and resistant cells. Kaznatcheev et al. [20] showed that alectinib and fibroblasts can change eco-evolutionary dynamics of NSCLC cells. Pressley, in Chapter 6 of her dissertation, examined adaptive therapy in an NSCLC model and showed that limited density dependence and variable evolvability can restrict the benefit of adaptive therapy [32]. These results suggest that evolutionary therapy depends not only on the presence of resistant cells, but also on how ecological context and resistance dynamics shape their competitive advantage.

NSCLC provides a useful setting for studying these issues because therapy resistance is common and the tumor microenvironment can strongly influence treatment response. Kaznatcheev et al. developed an experimental game assay using alectinib-sensitive and alectinib-resistant H3122 NSCLC cells grown under four conditions: with or without alectinib and with or without cancer-associated fibroblasts (CAFs) [20]. Their work showed that alectinib and CAFs can switch the evolutionary game played by sensitive and resistant cells, emphasizing that therapy and microenvironmental context alter cell-cell interactions. The same dataset has since been reanalyzed with different modeling approaches. Soboleva et al. used a polymorphic Gompertzian model to show that heterogeneous cancer population models can capture *in-vitro* and *in-vivo* treatment response dynamics [25]. Garjani et al. then compared population dynamic models that include density dependence, frequency-dependent competition, and different drug efficacy terms, showing that the inferred interaction dynamics depend on the chosen biological structure of the model [33]. Together, these studies demonstrate the richness of the dataset and the importance of model assumptions. However, they largely treat resistance as a fixed phenotype or focus on ecological interactions without asking whether the resistance trait itself changes during the experiment.

Here, we reuse this previously studied NSCLC dataset with a different aim. Rather than treating reanalysis of an existing dataset as a limitation, we use it as an opportunity to test a new biological hypothesis on a well-characterized experimental system: whether pre-existing resistant cells should be modeled as a subpopulation with a resistance trait that can evolve during treatment. This allows us to ask what earlier analyses could not directly address: whether the fitness effect of resistance is constant, or whether it depends on initial tumor composition and ecological competition between sensitive and resistant cells. In this sense, the dataset is not merely reused; it is reinterpreted through a model designed to reveal a different layer of resistance biology.

In this paper, we study how evolving resistance affects treatment outcomes in NSCLC. We use a polymorphic game-theoretic model in which sensitive and resistant cells compete ecologically, while resistance in the resistant population evolves as a continuous trait that affects both resistant-cell growth and treatment efficacy. The model assumes that resistance may pre-exist treatment, but that the resistant phenotype is not fixed. This formulation yields four possible regimes: resistance may carry a fitness cost or confer a growth advantage, and increasing resistance may either reduce or increase treatment efficacy. We fit the model to longitudinal *in-vitro* data from sensitive and resistant NSCLC cells seeded at different initial proportions and grown with or without alectinib and fibroblasts [20]. We then analyze how initial population composition influences inferred resistance dynamics and the feasibility of stabilization therapy, and we use dose simulations to examine how constant treatment dose affects time to progression. By linking pre-existing resistance, evolving resistance traits, and ecological competition, our work provides a framework for identifying when evolutionary therapy principles are likely to succeed or fail in NSCLC.

## 2 Materials and Methods

### 2.1 Background

We analyzed longitudinal *in-vitro* data originally generated by Kaznatcheev et al. [20]. The dataset contains treatment-sensitive parental H3122 NSCLC cells and an alectinib-resistant derivative cell line. Cells were grown either as monocultures or mixed cultures under four environmental conditions: with or without alectinib and with or without cancer-associated fibroblasts (CAFs). Sensitive and resistant cells were differentially labeled with fluorescent markers, allowing their population sizes to be measured over time. In the mixed-culture experiments, the two cancer cell types were seeded at different initial proportions, making the dataset especially useful for studying how initial tumor composition and cell-cell competition influence treatment response.

The same dataset has been used in subsequent modeling studies. Soboleva et al. fitted a polymorphic Gompertzian model to this *in-vitro* data and showed that this model can capture heterogeneous cancer population dynamics under treatment [25]. Their analysis also connected the *in-vitro* dataset to broader questions about adaptive therapy and tumor relapse in *in-vivo* data. However, they did not explicitly model direct competition coefficients or an evolving resistance trait. Garjani et al. later used the dataset to compare multiple population dynamic models, including logistic, Gompertzian, and von Bertalanffy growth, together with different drug efficacy terms [33]. Their analysis explicitly incorporated density dependence, frequency-dependent competition, and treatment response, showing that CAFs and alectinib can alter inferred competition and coexistence between sensitive and resistant cells. However, they treated resistance as a fixed trait.

### 2.2 Mathematical model

We use a polymorphic game-theoretic model to capture eco-evolutionary dynamics of the NSCLC cells under treatment, proposed in our earlier work [27]. This model assumes two cancer cell populations, treatment-sensitive (S) and treatment-resistant (R) ones. Let *x*_*i*_(*t*) ∈ ℝ_+_ and 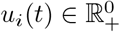 define the population and evolving resistance of all cells of type *i* ∈ *{S, R}* at time *t*, and **x**(*t*) = (*x*_*i*_(*t*))_*i*∈*{S,R}*_ and **u**(*t*) = (*u*_*i*_(*t*))_*i*∈*{S,R}*_ . We assume that only resistant cells have the capacity to evolve resistance 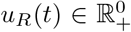 and sensitive cells cannot evolve resistance, i.e., *u*_*S*_(*t*) = 0 for all *t*.

Ecological dynamics of cells of type *i* are defined as

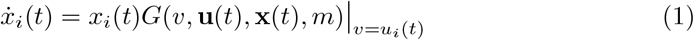

where *G* refers to the fitness-generating function from Darwinian dynamics [34] and *v* is the resistance of the focal cell. In addition, *m* ≥ 0 corresponds to a treatment dose applied by a physician.

We assume that the treatment resistance *u*_*R*_(*t*), may evolve in the direction of the fitness gradient 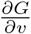:

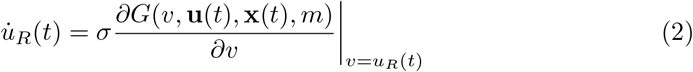

The parameter *σ >* 0 scales the speed at which the resistance changes.

The G-function is defined as follows:

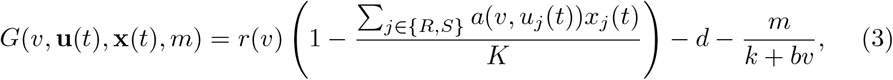

thus assuming a logistic cancer growth model with carrying capacity *K*. In (3), *a*(*u*_*i*_, *u*_*j*_) = *α*_*ij*_ defines the competition effect of cancer cells of type *j* on those of type *i*, with *i*, ∈ {*j R, S}*. The term *r*(*v*) is the growth rate of the focal cancer cell type, depending on their resistance rate *v*, defined as *r*(*v*) = *r*_max_ *e*^−*g v*^, representing cost of resistance if *g >* 0 and gain of resistance if *g <* 0. The term *d* represents the natural death rate [35], while the term 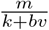 represents the Michaelis-Menten treatment-induced death rate [36], where *k >* 0 represents innate drug resistance and *b* determines how the evolving resistance trait changes treatment efficacy. If *b >* 0, increasing resistance reduces treatment-induced mortality; if *b <* 0, increasing resistance increases treatment-induced mortality.

In the remainder of this paper, we drop the time symbol *t* for the sake of simplicity and readability of our notation.

The full model (1)–(3) of cancer eco-evolutionary dynamics then reads as follows:

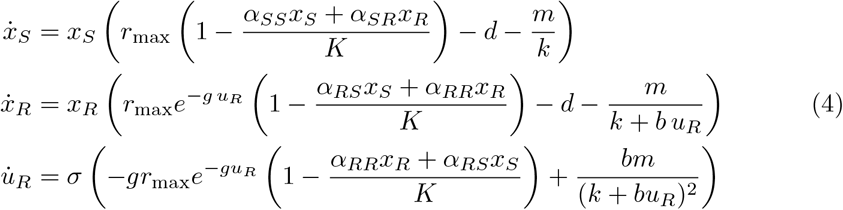

### 2.3 Fitting procedure

We fitted the model to each well separately using nonlinear constrained optimization in MATLAB (MathWorks, RRID:SCR 001622). For each well, the model was initialized using the experimentally measured initial sensitive and resistant cell populations. Parameter values were then estimated by minimizing the mean squared error between the measured and simulated sensitive and resistant cell populations over time. This procedure yielded one fitted parameter set per well.

After fitting, we first compared the distributions of fitted parameters across the four experimental conditions defined by the presence or absence of alectinib and fibroblasts. We then examined whether additional experimental features explained variation in the fitted parameters. The initial composition of sensitive and resistant cells provided the clearest separation of fitted parameter patterns. Therefore, in addition to the four resistance regimes defined by the signs of *g* and *b*, we classified wells according to whether sensitive or resistant cells were initially in the majority.

Examples of fitted trajectories are presented in Appendix D. The initial seeding structure of the wells is provided in Appendix C. For example, well (8, 5) denotes the well in the 8th row and 5th column, where the initial population consisted of 97.5% sensitive cells and 2.5% resistant cells. In this well, both alectinib and fibroblasts were present.

### 2.4 Analytical and simulation-based analysis

We analyzed the model to determine how evolving resistance affects the existence and stability of tumor equilibria under treatment. The sign of *g* determines whether increasing resistance reduces or increases the intrinsic growth rate of resistant cells. We refer to *g >* 0 as a cost of resistance and *g <* 0 as a gain of resistance. The sign of *b* determines whether increasing resistance decreases or increases treatment-induced mortality. Together, the signs of *g* and *b* define four resistance regimes.

For each regime, we examined the existence and local stability of equilibria corresponding to tumor elimination, fully sensitive tumors, fully resistant tumors, and coexistence of sensitive and resistant cells. Because the fitted data showed no cases with negative resistance benefit, the main analytical focus was on the case *b >* 0, in which increasing resistance reduces treatment efficacy. We paid particular attention to the gain-of-resistance case, *g <* 0, because in this regime resistance evolution can increase resistant-cell growth while simultaneously reducing treatment efficacy.

To study the feasibility of stabilization therapy, we analyzed whether stable tumor equilibria can be maintained at a bounded tumor burden under constant treatment dose. In the gain-of-resistance case, we derived conditions under which stable equilibria can exist and assessed whether such equilibria include resistant cells or require resistant-cell exclusion. Details of the equilibrium calculations and stability analysis are provided in Appendix A.

We also used numerical simulations to examine how constant treatment dose affects time to progression. Total tumor burden was defined as *X*(*t*) = *x*_*S*_(*t*) + *x*_*R*_(*t*), and time to progression was defined as the time at which *X*(*t*) reached 70% of the carrying capacity. For selected parameter sets, we simulated the model over a range of constant doses and compared the resulting time to progression with the initial tumor growth rate, 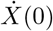. This allowed us to identify cases in which maximum tolerated dosing maximized time to progression and cases in which intermediate dosing performed better.

### 2.5 Reproducibility and Rigor

Because this study relies on mathematical modeling of a previously published invitro dataset using H3122 NSCLC cells (RRID:CVCL 5160), criteria regarding a priori power analysis, randomization, and investigator blinding were not applicable. Sex as a biological variable was not considered as the study utilizes an established cell line. The MATLAB (MathWorks, RRID:SCR 001622) code used for the nonlinear constrained optimization is available on GitHub : https://github.com/Mohammad2aAq/NSCLC-optimization-model. The original dataset analyzed in this study is available at https://github.com/kaznatcheev/GameAssay.

## 3 Results

### 3.1 When resistant cells are initially rare, evolving resistance is associated with a growth advantage

Consistent with the effects of initial population composition on evolutionary speed and innate resistance, we next examined how the initial proportion of resistant cells influenced the inferred fitness effect of evolving resistance. This effect is determined by the parameter *g*: when *g >* 0, increasing resistance reduces the intrinsic growth rate of resistant cells, corresponding to a cost of resistance; when *g <* 0, increasing resistance increases the intrinsic growth rate of resistant cells, corresponding to a gain of resistance.

After fitting the model to the data, we found that the sign of *g* depended strongly on the initial composition of sensitive and resistant cells under treatment. When resistant cells were initially in the minority, the fitted values of *g* were predominantly negative, indicating that evolving resistance was associated with a growth advantage. In contrast, when resistant cells were initially in the majority, the fitted values of *g* were predominantly positive, indicating a cost of resistance. The distributions of fitted values of *g* are shown in Figure 1.

**Fig. 1:**
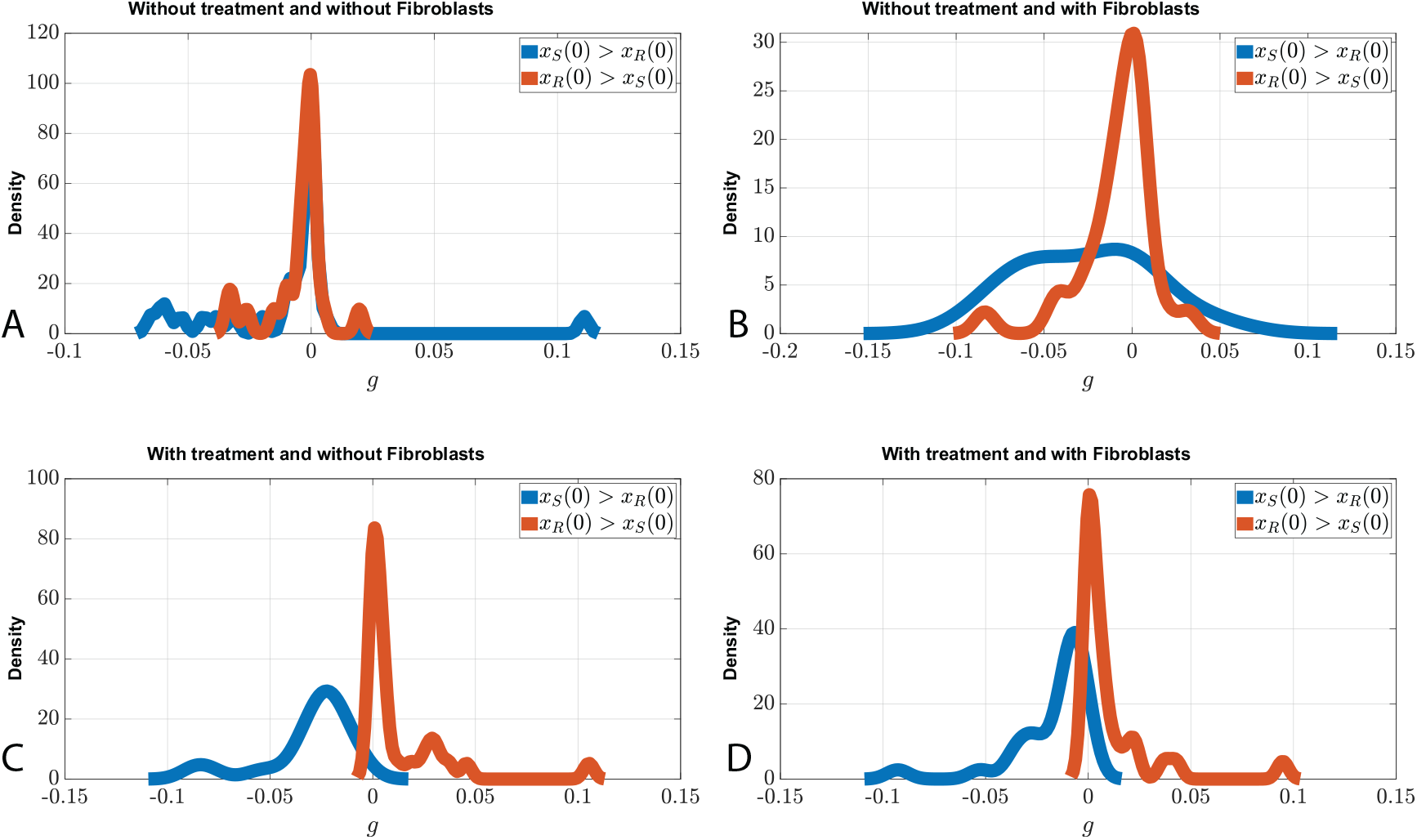
Kernel density estimation plots for the inferred cost or gain of resistance under four experimental conditions: with or without alectinib, and with or without fibroblasts. Red curves indicate wells in which resistant cells outnumbered sensitive cells at the onset of the experiment. Blue curves indicate wells in which sensitive cells outnumbered resistant cells at the onset of the experiment. In A and B, where alectinib is absent, the red and blue distributions are similar. In C and D, where alectinib is present, fitted values of *g* are predominantly positive when resistant cells are initially more abundant and predominantly negative when sensitive cells are initially more abundant. Positive values of *g* indicate a cost of resistance, whereas negative values indicate a gain of resistance.

Thus, under alectinib, the same resistance trait was inferred to have different fitness effects depending on the initial ecological context: a growth advantage when resistant cells were initially rare and a fitness cost when resistant cells were initially common.

### 3.2 Initial population composition affects the inferred speed of resistance evolution

After fitting the NSCLC dataset to the Darwinian dynamics model (4), we found that the inferred speed of resistance evolution depended on the initial composition of sensitive and resistant cells. When sensitive cells were initially in the majority, the estimated evolutionary speed of the resistant-cell resistance trait was higher than when resistant cells were initially in the majority. Across wells with and without fibroblasts, the mean estimated evolutionary speed was 133.59 when sensitive cells were initially more abundant and 65.54 when resistant cells were initially more abundant. In the presence of both alectinib and fibroblasts, the corresponding mean values were 150.90 and 45.04, respectively.

The distribution of estimated evolutionary speeds is shown in Figure 2. In the presence of alectinib, wells with initially more sensitive cells showed higher estimated evolutionary speeds than wells with initially more resistant cells. In contrast, in the absence of alectinib, the distributions were more similar. The presence or absence of fibroblasts did not substantially alter these patterns.

**Fig. 2:**
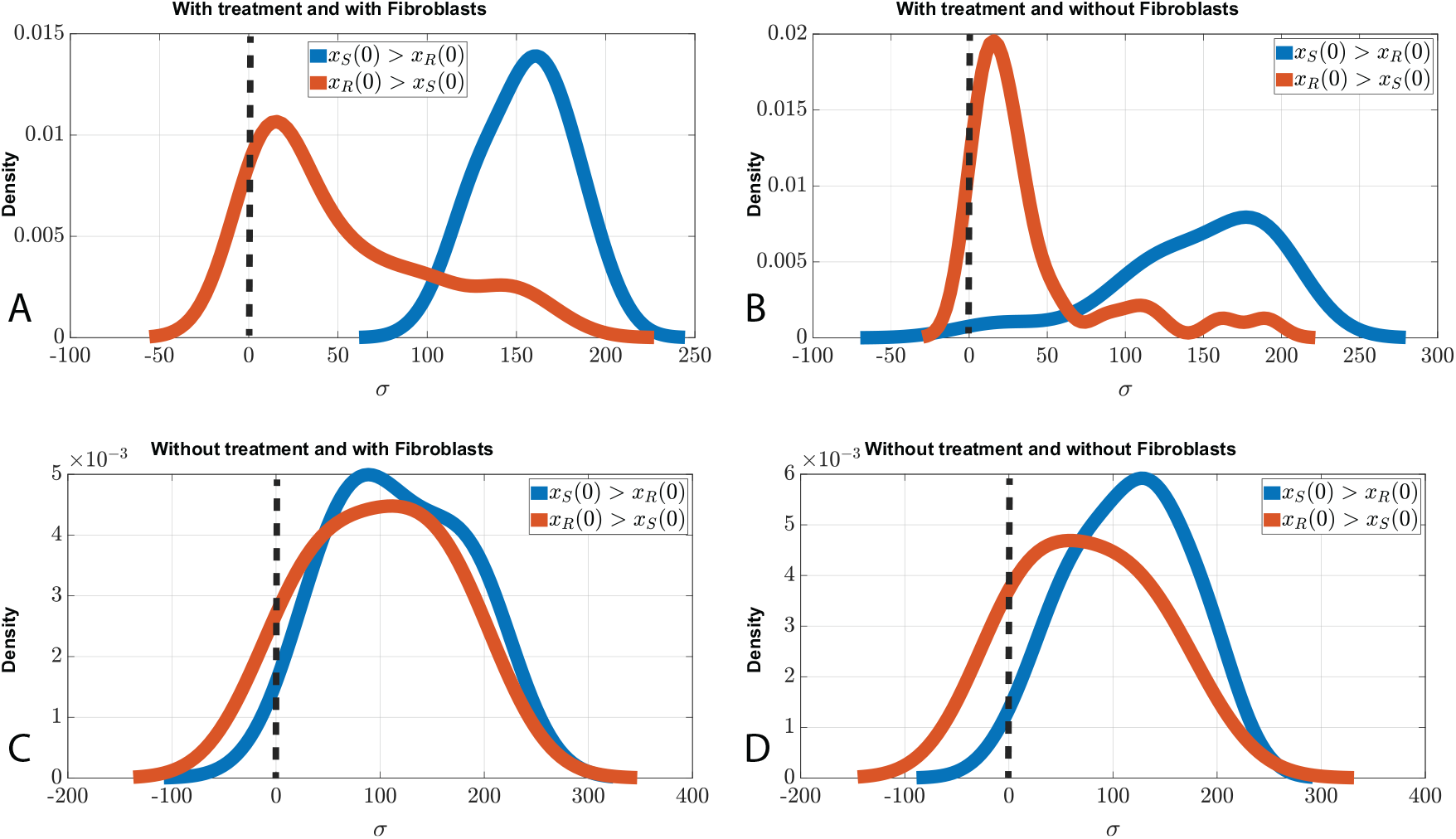
Kernel density estimation plots for the inferred evolutionary speed of the resistant-cell resistance trait under four experimental conditions: with or without alectinib, and with or without fibroblasts. Red curves indicate wells in which resistant cells outnumbered sensitive cells at the onset of the experiment. Blue curves indicate wells in which sensitive cells outnumbered resistant cells at the onset of the experiment. In A and B, where alectinib is present, wells with initially more sensitive cells show higher estimated evolutionary speeds. In C and D, where alectinib is absent, the red and blue distributions are more similar. All fitted evolutionary speed values are positive; the small negative tails arise from kernel density estimation.

### 3.3 Initial population composition affects inferred innate resistance

We also found that the inferred innate resistance parameter, *k*, depended on the initial composition of sensitive and resistant cells. In the model (4), *k* modulates treatment-induced mortality in both sensitive and resistant cells: higher values of *k* correspond to a weaker treatment effect. In the presence of alectinib, wells in which sensitive cells were initially in the majority had higher inferred innate resistance than wells in which resistant cells were initially in the majority. The mean estimated value of *k* was 318.73 when sensitive cells were initially more abundant and 56.67 when resistant cells were initially more abundant.

The distribution of estimated innate resistance values is shown in Figure 3. Both with and without fibroblasts, wells with initially more sensitive cells showed higher inferred innate resistance. Thus, in the fitted model, cultures with fewer resistant cells at the onset of treatment were associated with a weaker inferred treatment effect.

**Fig. 3:**
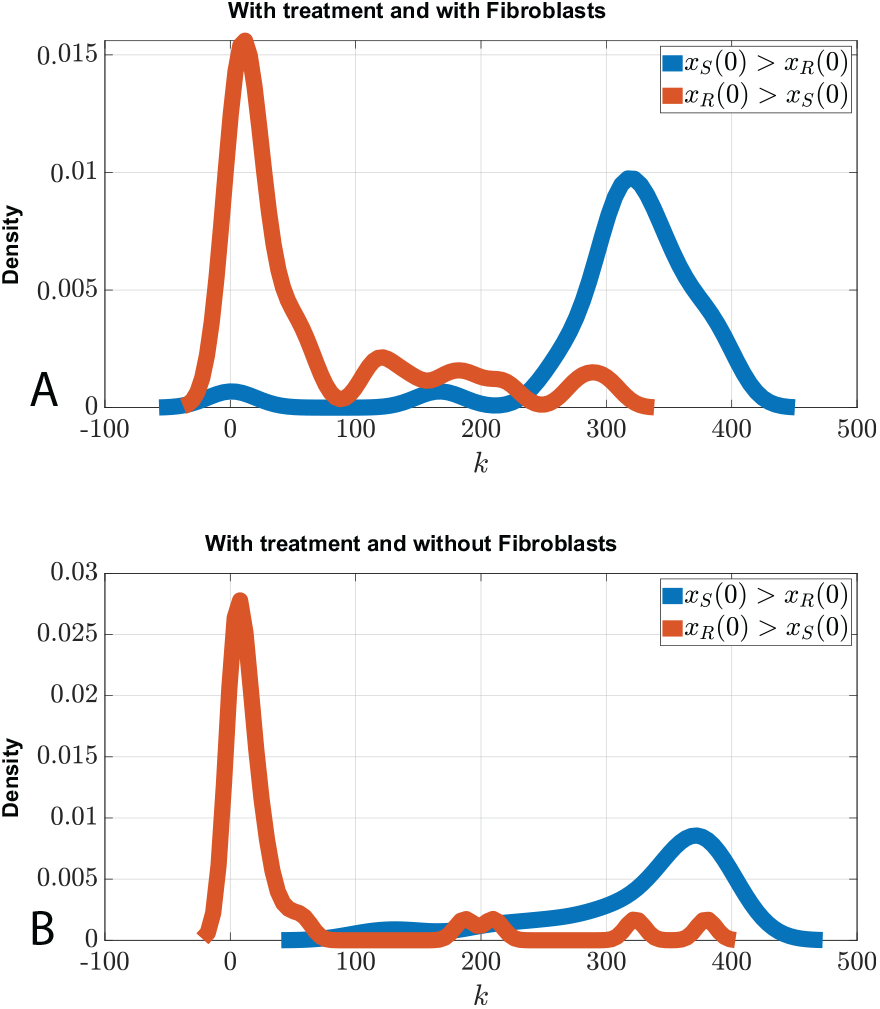
Kernel density estimation plots for the inferred innate resistance parameter, *k*, in the presence of alectinib, with or without fibroblasts. Red curves indicate wells in which resistant cells outnumbered sensitive cells at the onset of the experiment. Blue curves indicate wells in which sensitive cells outnumbered resistant cells at the onset of the experiment. In A, fibroblasts are present, while in B, fibroblasts are absent. In both cases, wells with initially more sensitive cells show higher inferred innate resistance.

### 3.4 In the gain-of-resistance regime, stabilization therapy requires resistant-cell exclusion

In stabilization therapy, treatment is chosen to maintain tumor burden at a stable, clinically tolerable level rather than to eradicate the tumor. In the setting considered here, this means applying a constant treatment dose that drives the cancer dynamics (4) toward a stable equilibrium with total tumor burden below a predefined threshold [28].

In the gain-of-resistance regime, however, such stabilization cannot maintain a tumor containing resistant cells. If a resistant population persists, its resistance trait grows without bound, and the system cannot converge to a fixed tumor equilibrium with resistant cells present. Consequently, in this regime, stabilization therapy can only maintain a stable tumor burden if resistant cells are excluded and the resulting tumor is fully sensitive. The analytical proof is presented in Appendix A. In addition, we showed in [27] that, when resistance carries a fitness cost, stabilization therapy can result in four different outcomes: a fully sensitive tumor, a fully resistant tumor, coexistence of resistant and sensitive cells, or tumor elimination.

For a stable fully sensitive equilibrium to exist, sensitive cells must be sufficiently competitive to exclude resistant cells. Therefore, when sensitive cells constitute the majority at the onset of treatment, stabilization therapy is more likely to drive the system toward a fully sensitive equilibrium in the gain-of-resistance regime. Because the presence of sensitive cells is required for competitive exclusion of resistant cells, this outcome is expected to be more feasible under lower treatment doses that preserve a sufficiently large sensitive-cell population. For instance, in wells (8, 1) and (8, 7), where sensitive cells initially constituted 97.5% and 80% of the population, respectively, the fitted dynamics indicate a fully sensitive equilibrium under lower doses of alectinib.

### 3.5 No NSCLC culture behaved as a Darwinian demon

A Darwinian demon is a hypothetical type that simultaneously improves multiple fitness-related traits without trade-offs [37]. In the present context, a Darwinian-demon-like regime would correspond to resistant NSCLC cells for which increasing resistance both increases intrinsic growth and undermines treatment control.

In the model (4), the gain-of-resistance condition is *g <* 0, meaning that increasing resistance increases the intrinsic growth rate of resistant cells. The second condition concerns the effect of evolving resistance on treatment efficacy, determined by *b*. When *b >* 0, increasing resistance reduces treatment efficacy, whereas *b <* 0 would correspond to a negative resistance benefit. The analytical characterization of this case is presented in Appendix B.

After fitting the model to the NSCLC data, we found no parameter set with *b <* 0. Thus, although some fitted cultures showed a gain of resistance (*g <* 0), none showed a Darwinian-demon regime. In all fitted cases, *b >* 0, meaning that treatment continued to reduce resistant-cell growth, although its efficacy declined as the resistance trait increased.

### 3.6 Initial experimental conditions are associated with fitted model parameters

We used linear models to assess how the fitted parameter values depended on the initial experimental conditions. Specifically, we used the initial number of resistant cells, *x*_*R*_(0), the presence or absence of alectinib, and the presence or absence of fibroblasts as predictors of the fitted parameters in the cancer dynamics model (4). For each fitted parameter, we evaluated the direction and significance of the corresponding regression coefficients. To correct for multiple testing, we used Benjamini-Hochberg false discovery rate (FDR) corrected p-values [38].

Figure 4 shows the associations between initial experimental conditions and fitted model parameters. Values above 2 correspond to FDR-corrected p-values below 0.01 and were considered strongly significant. The final column shows the FDR-corrected p-value for the overall linear model for each fitted parameter.

**Fig. 4:**
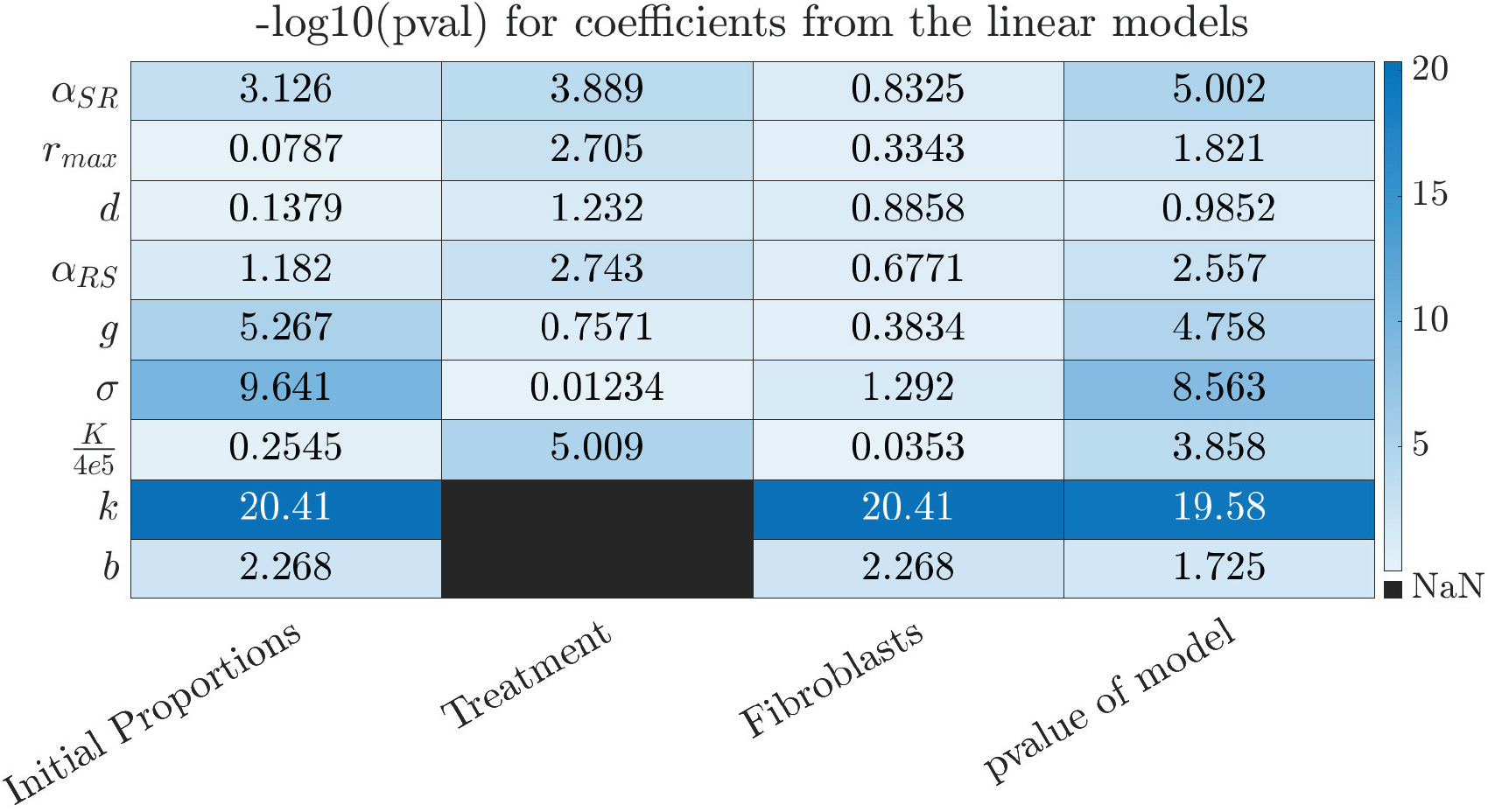
Associations between initial experimental conditions and fitted parameters of the cancer dynamics model (4). The heatmap shows −log_10_ transformed Benjamini-Hochberg FDR-corrected p-values for linear models using the initial number of resistant cells, *x*_*R*_(0), the presence of alectinib, and the presence of fibroblasts as predictors. Values above 2 correspond to FDR-corrected p-values below 0.01. The final column shows the FDR-corrected p-value for the overall linear model.

The initial number of resistant cells, *x*_*R*_(0), was significantly associated with several fitted parameters, including the competition coefficient *α*_*SR*_, the cost or gain of resistance *g*, the evolutionary speed *σ*, the innate resistance parameter *k*, and the resistance benefit parameter *b*. Alectinib treatment was significantly associated with the competition coefficients *α*_*SR*_ and *α*_*RS*_, the maximum growth rate *r*_max_, and the carrying capacity *K*. Fibroblasts were significantly associated with the innate resistance parameter *k* and the resistance benefit parameter *b*. Among these associations, *g* and *b* were the parameters most specifically linked to the initial number of resistant cells, supporting the conclusion that initial population composition is particularly informative about the inferred fitness effect and treatment benefit of evolving resistance.

Because these linear models did not account for interactions between initial experimental conditions, we fitted a second set of linear models that included interaction terms between the initial number of resistant cells, alectinib treatment, and fibroblasts. The corresponding FDR-corrected p-values are shown in Figure 5.

**Fig. 5:**
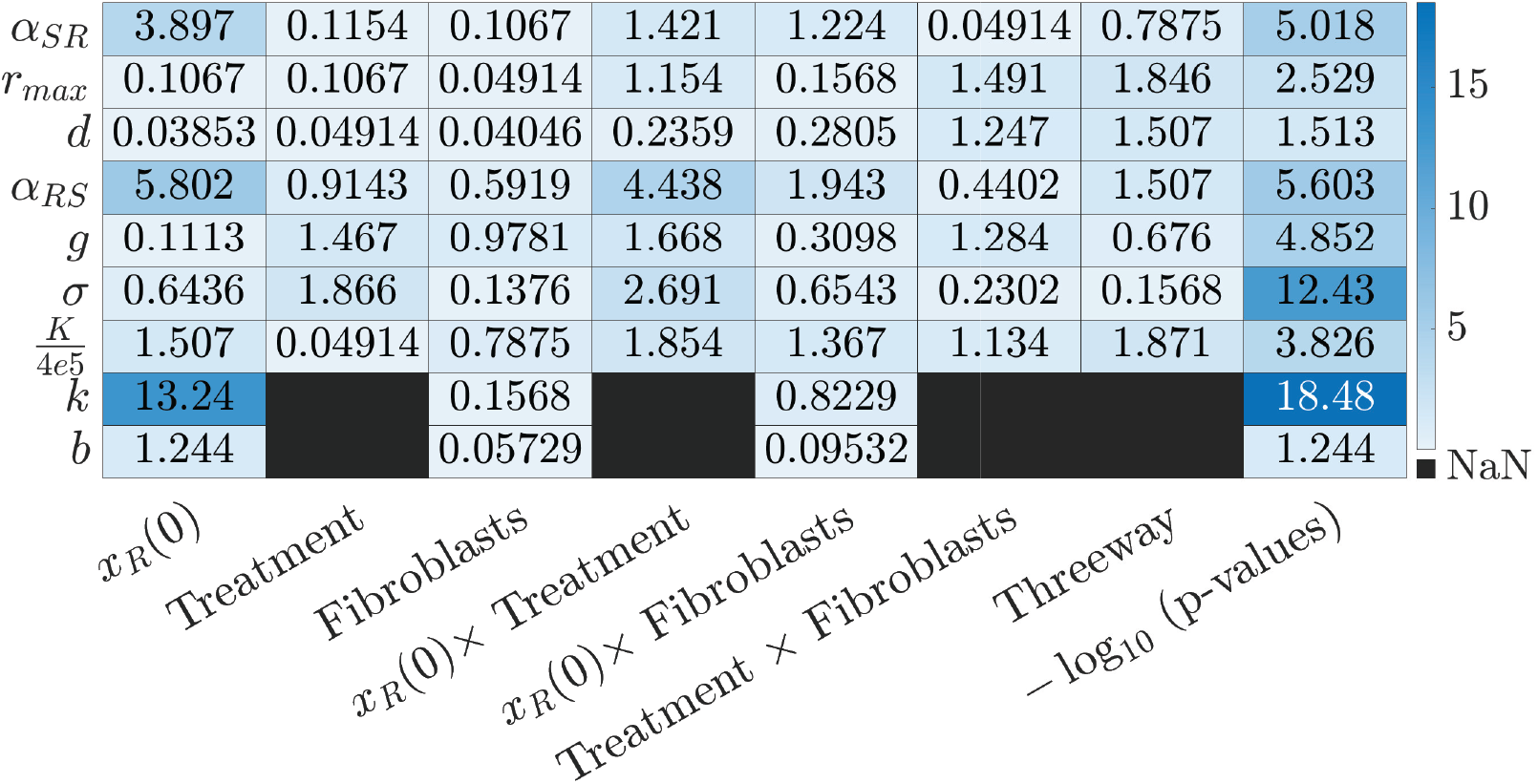
Associations between interaction terms among initial experimental conditions and fitted parameters of the cancer dynamics model (4). The heatmap shows −log_10_ transformed Benjamini-Hochberg FDR-corrected p-values for interaction terms between the initial number of resistant cells, alectinib treatment, and fibroblasts.

Using the same significance threshold, the only interaction term with a strong association was the interaction between the initial number of resistant cells and alectinib treatment. This interaction was significant for *α*_*RS*_ and *σ*, suggesting that the effect of initial population composition on these parameters depends on whether treatment is present.

### 3.7 Correlations among fitted model parameters

To identify associations among fitted parameters of the cancer dynamics model (4), we calculated Spearman rank correlations for each pair of parameters across wells. The results are shown in Figure 6. For readability, the main-text figure omits the numerical correlation values; larger versions including these values are provided in Appendix E.

**Fig. 6:**
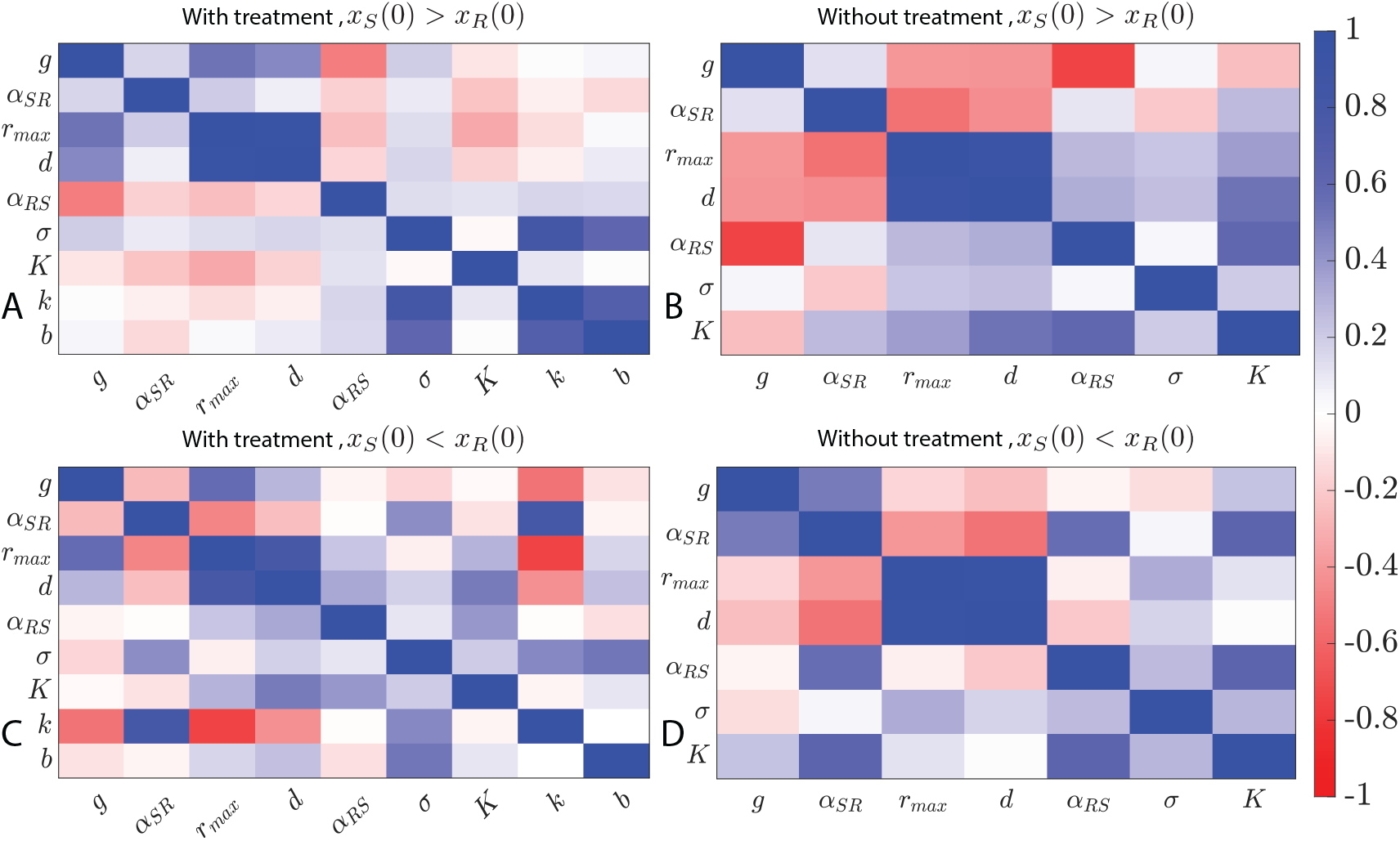
Spearman rank correlations among fitted parameters of the cancer dynamics model (4) under four experimental scenarios. Numerical correlation values are omitted here for readability and are provided in Appendix E.

The correlation structure differed across experimental scenarios, but one pattern was consistent: the maximum growth rate, *r*_max_, and the natural death rate, *d*, were strongly positively correlated in all scenarios. The Spearman rank correlation between these two parameters exceeded 0.9 in all cases except for the treated scenario in which resistant cells were initially in the majority, where the correlation was 0.79. This relationship is also visible in the scatter plots shown in Figure 7.

**Fig. 7:**
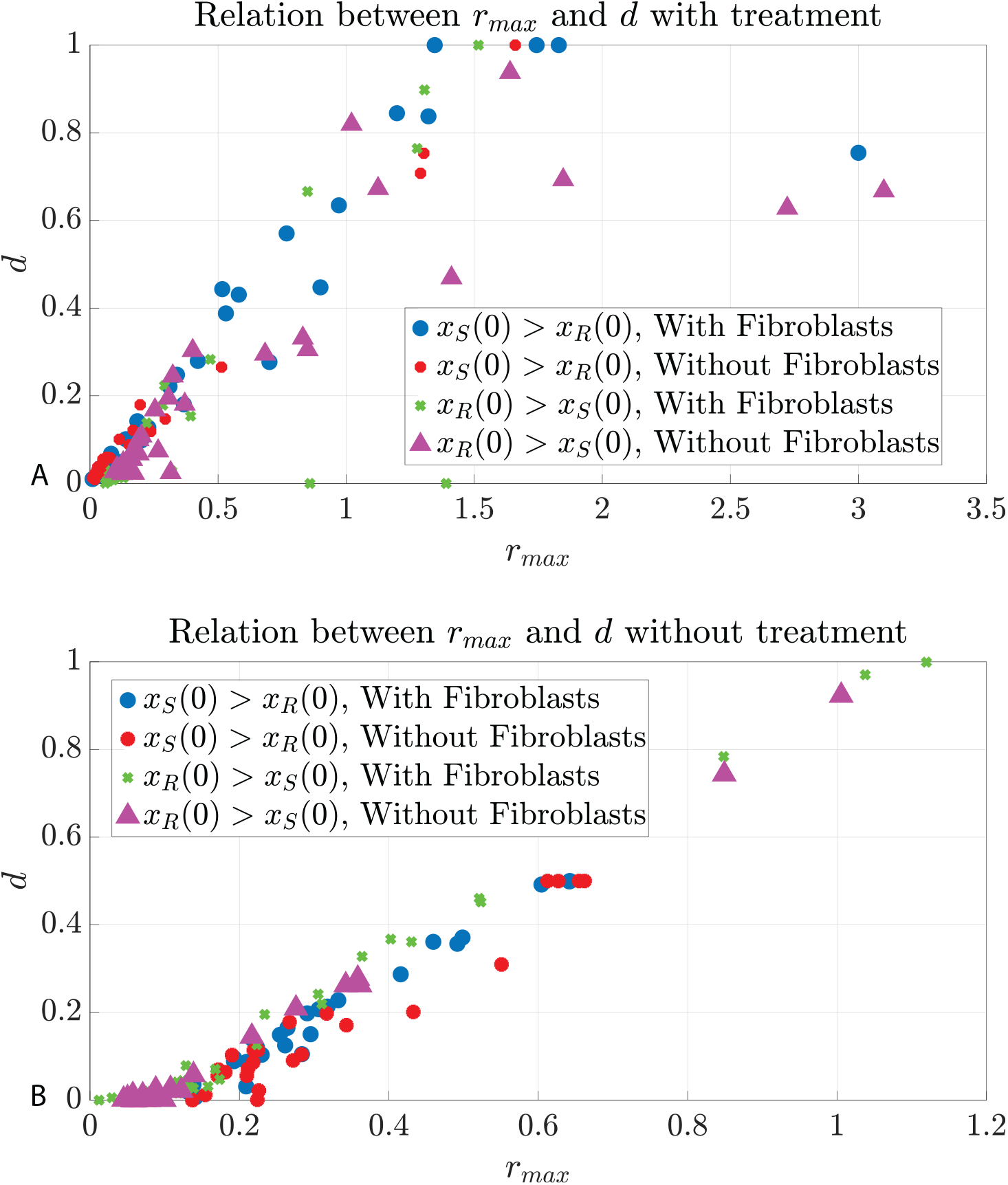
Relationship between the maximum growth rate, *r*_max_, and the natural death rate, *d*, across fitted wells. These parameters are strongly positively correlated across experimental scenarios. A: wells treated with alectinib. B: wells without alectinib.

Additional parameter associations depended on the experimental context. In the presence of alectinib, when resistant cells were initially in the majority, *r*_max_ and innate resistance *k* were negatively correlated, whereas the competition coefficient *α*_*SR*_ and innate resistance *k* were positively correlated. In contrast, when sensitive cells were initially in the majority under alectinib treatment, the evolutionary speed *σ* and innate resistance *k* were positively correlated. Scatter plots for these additional associations are provided in Appendix E.

### 3.8 Treatment dose affects time to progression in the gain-of-resistance regime

In the gain-of-resistance regime, *g <* 0, we showed that stabilization therapy can maintain a stable tumor burden only if resistant cells are excluded eventually, resulting in a fully sensitive tumor. When such a stable, clinically tolerable equilibrium exists, the appropriate constant treatment dose is the one that drives the system toward this equilibrium, as discussed in Subsection 3.4. If no such stable tumor burden exists, treatment may instead aim to delay progression. Here, we define time to progression (TTP) as the time at which the total tumor burden reaches 70% of its carrying capacity [12].

Let *X*(*t*) = *x*_*S*_(*t*) + *x*_*R*_(*t*) denote the total tumor burden. From the evolutionary equation in (4), when *g <* 0 the resistance trait *u*_*R*_ is strictly increasing. Moreover,

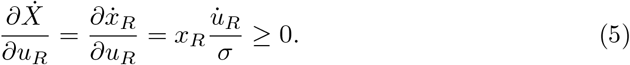

Thus, as the resistance trait increases, the rate of changes in the total tumor burden also increases. This conclusion holds independently of the treatment dose. We therefore considered how different constant treatment doses affect TTP in this regime.

Two cases can be distinguished.

#### Case I: when any treatment dose *m* ∈ [0, 1] produces initial tumor growth, the maximum tolerated dose maximizes time to progression

In this case, since 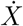 increases as *u*_*R*_ increases, the tumor burden continues to grow and no constant treatment dose can stabilize the tumor. However,

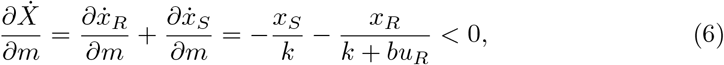

thus, higher treatment doses reduce the initial rate of tumor growth. In this case, the maximum tolerable dose slows tumor growth and can increase TTP. This situation is illustrated in Figure 8A.

**Fig. 8:**
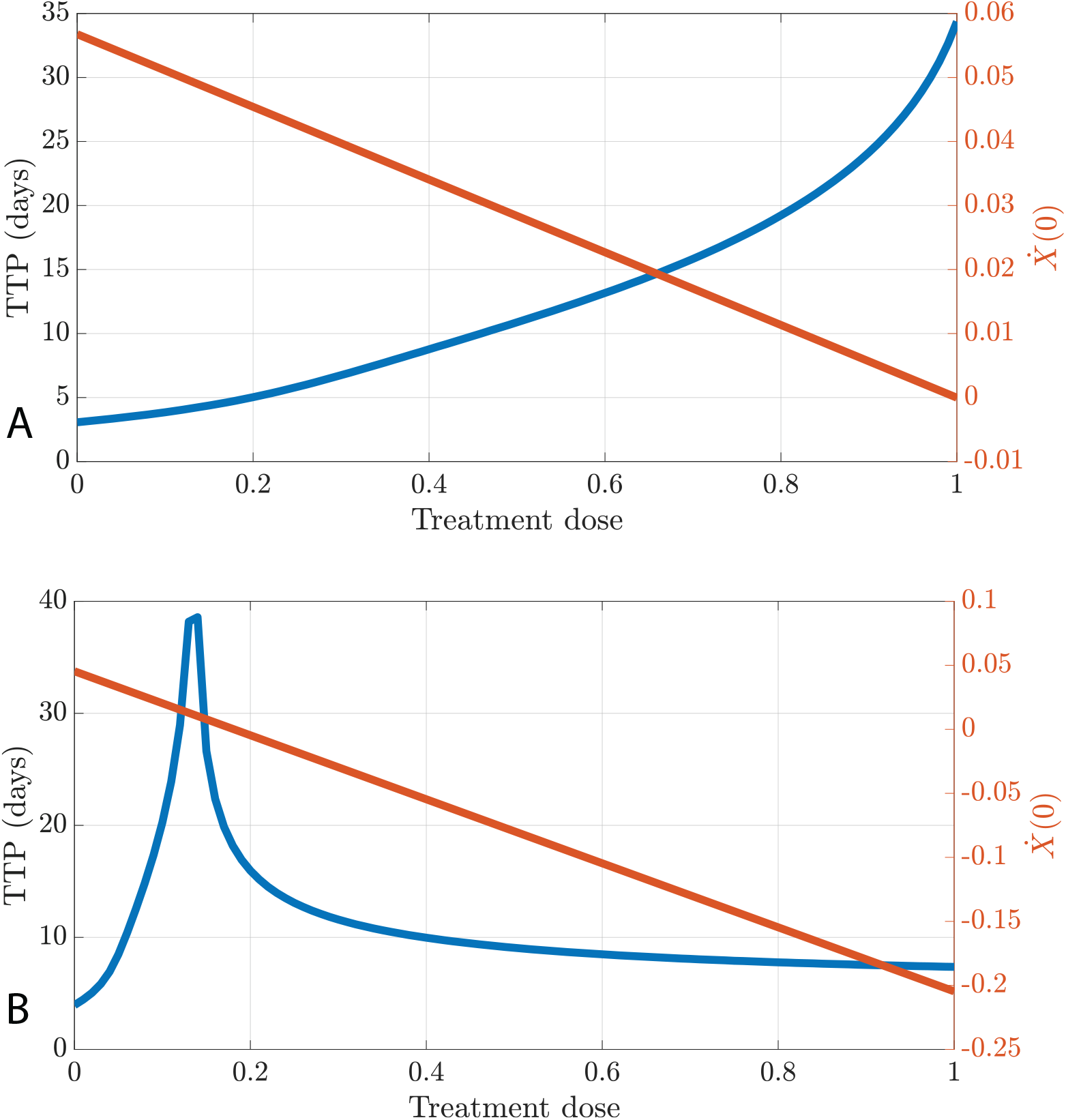
Effect of constant treatment dose on time to progression (TTP) in the gain-of-resistance regime. In both panels, the blue curve shows TTP, and the red curve shows the initial rate of change in total tumor burden, 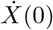. A: The tumor burden initially increases for all treatment doses. In this case, the maximum tolerated dose gives the longest TTP. Parameters: *g* = −0.03, *K* = 10000, *r*_max_ = 0.55, *k* = 10, *b* = 12, *d* = 0.01, *σ* = 10, *α*_*SS*_ = 1, *α*_*RR*_ = 1, *α*_*SR*_ = 1.5, *α*_*RS*_ = 1.9. B: The initial rate of change in tumor burden varies from positive to negative across treatment doses. In this case, an intermediate dose, 21% of the maximum tolerated dose, gives the longest TTP and corresponds to 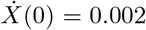. Parameters: *g* = −0.03, *K* = 10000, *r*_max_ = 0.35, *k* = 2, *b* = 10, *d* = 0.01, *σ* = 10, *α*_*SS*_ = 1, *α*_*RR*_ = 1, *α*_*SR*_ = 1.5, *α*_*RS*_ = 1.9.

#### Case II: when an intermediate dose keeps initial tumor growth near zero, this dose can outperform the maximum tolerated dose in terms of TTP

In this case, there exists a treatment dose *m* for which the initial rate of change in tumor burden is zero. By (6), doses above *m* produce an initial decrease, whereas doses below *m* produce an initial increase in tumor burden. However, a larger initial reduction in tumor burden can also intensify competitive release of the resistant population, allowing the tumor to reach the progression threshold earlier. Therefore, the dose that maximizes TTP need not be the maximum tolerated dose. Instead, intermediate doses that keep 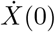 close to zero can produce a higher TTP. This situation is illustrated in Figure 8B.

##### Remark 1.

*Because this argument is based on the initial rate of change in total tumor burden*, 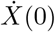, *it does not imply that the dose satisfying* 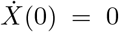 *always maximizes TTP. Later dynamics also matter. Nevertheless, in the gain-of-resistance regime, doses that keep the initial tumor growth rate close to zero provide a useful rule of thumb for identifying treatments that may prolong TTP*.

In the cost-of-resistance regime, *g >* 0, stable equilibria may include a fully sensitive tumor, a fully resistant tumor, a mixed sensitive-resistant tumor, or tumor elimination, as shown in [27]. In that regime, the appropriate constant dose is the dose associated with the desired stable equilibrium and clinically tolerable tumor burden.

### 3.9 Within the gain-of-resistance regime, the patients with lower gains reach the stabilization burden faster

Within the gain-of-resistance regime (*g <* 0), stabilization therapy can maintain a clinically acceptable tumor burden only if resistant cells are excluded and the cancer dynamic converges to a fully sensitive equilibrium, as discussed in Appendix A. The feasibility of this outcome depends on whether the conditions for existence and stability of the fully sensitive equilibrium can be satisfied.

According to (11), the gain-of-resistance parameter *g* enters the condition for the existence of the fully sensitive equilibrium through the evolutionary equation for the resistant-cell resistance trait. Since *g <* 0, this condition requires

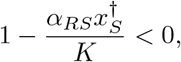

meaning that sensitive cells must be sufficiently competitive against resistant cells. Thus, stabilization in this regime is possible only when sensitive cells can exclude resistant cells and the resulting fully sensitive equilibrium lies below the predefined clinically acceptable tumor burden.

As illustrated in Figure 9, weaker gains of resistance are stabilized on a burden below 70% of the carrying capacity, faster than the higher gains. Moreover, for the gains higher than 0.07, the stabilization is not possible and tumor burden reaches close to the carrying capacity which is not tolerable for the patient.

**Fig. 9:**
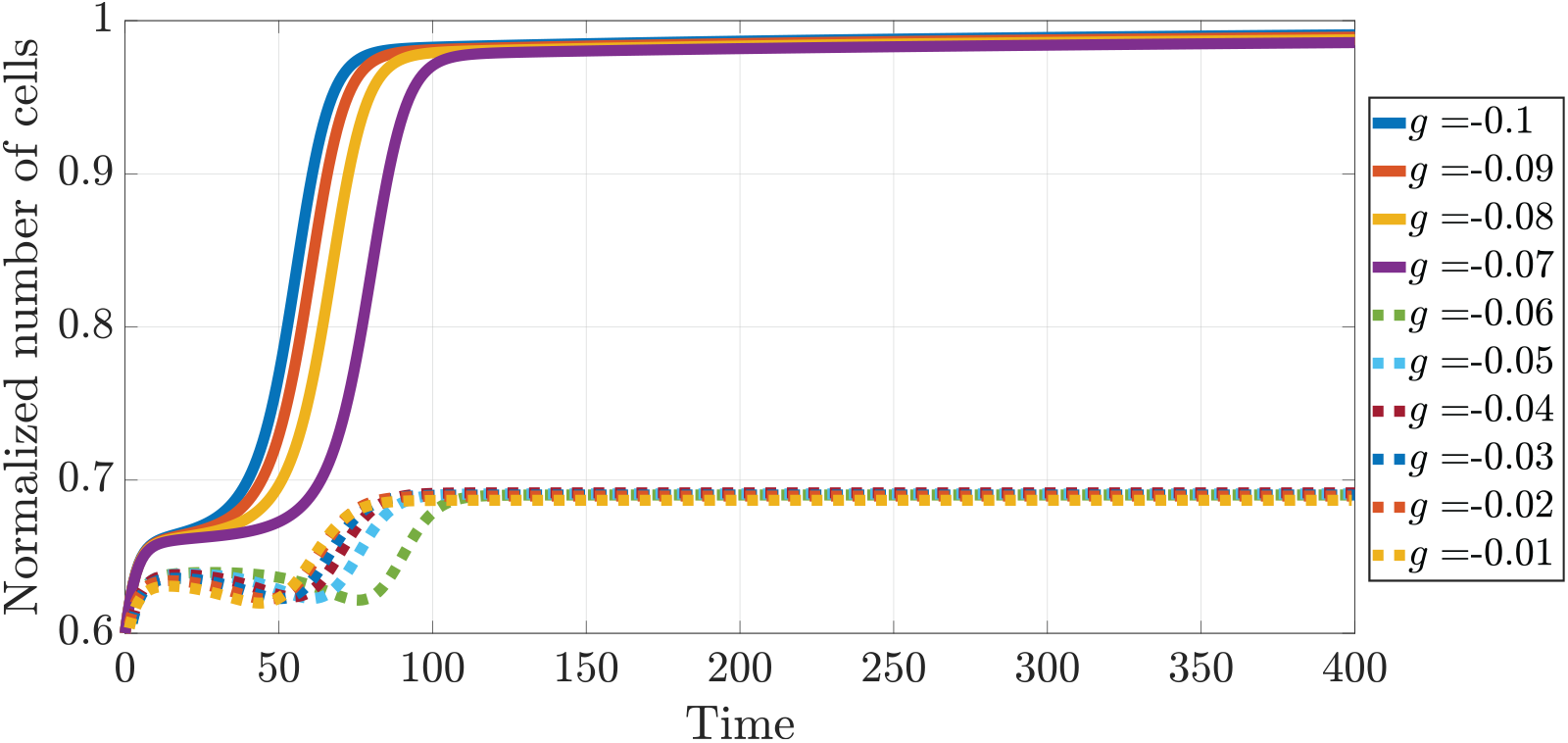
Effect of the gain-of-resistance parameter on stabilization outcomes in the gain-of-resistance regime. For the parameter values used in these simulations, weaker gains of resistance are associated with stabilization below 70% of the carrying capacity, whereas stronger gains of resistance are associated with higher tumor burdens.

In addition, from (11), we can see that for higher gains, the resistance rate at the equilibrium would be lower compared to the cases with lower gains. Thus, in the case of cancer relapse, patients with higher gains, would benefit from lower resistance rates.

In these examples of Figure 9, sensitive cells are more competitive than resistant cells, that is, *α*_*RS*_ *> α*_*SR*_. This competitive asymmetry is necessary for sensitive cells to exclude resistant cells and thereby enable stabilization at a fully sensitive equilibrium. Moreover, because the persistence of sensitive cells is essential for this outcome, stabilization therapy in the gain-of-resistance regime is more likely to succeed when treatment doses are low enough to preserve a sufficiently large sensitive-cell population.

## 4 Discussion

Evolutionary therapy is often motivated by the idea that treatment-sensitive and treatment-resistant cancer cells compete, and that this competition can be exploited to delay progression. A central assumption in many such models is that resistance is a fixed cellular state that carries a fitness cost in the absence of therapy. Here, we tested a more dynamic alternative: resistance was allowed to evolve as a continuous trait, and this trait could alter both the growth rate of resistant cells and their response to treatment. By fitting this model to longitudinal *in-vitro* data from mixed NSCLC cultures, we found that the inferred effect of resistance was not fixed. Instead, it depended strongly on the initial composition of sensitive and resistant cells.

The most important result is that resistance was associated with different fitness consequences in different ecological contexts. When resistant cells were initially rare, the fitted model inferred faster evolution of resistance and a gain-of-resistance regime, meaning that increasing resistance was associated with a higher intrinsic growth rate. When resistant cells were initially frequent, increasing resistance was instead associated with a fitness cost. Thus, resistance did not behave as a single, context-independent property of the resistant cell line. Rather, the same experimental system could support either cost-of-resistance or gain-of-resistance dynamics depending on the starting composition of the culture. This finding supports the broader view that treatment response is shaped not only by molecular resistance mechanisms, but also by ecological interactions among cancer cell populations and their environment [4, 12, 20].

This context dependence has direct implications for evolutionary therapy. Many evolutionary therapy strategies rely on maintaining treatment-sensitive cells to suppress resistant cells through competition [11–13, 39]. Such strategies are most straightforward when resistance carries a cost: reducing treatment can allow sensitive cells to recover and competitively restrain resistant cells. Our results show why this assumption needs to be tested rather than imposed. If increasing resistance also increases resistant-cell growth, then the ecological logic of containment changes. In the gain-of-resistance regime, our analytical results show that a stable tumor equilibrium containing resistant cells cannot be maintained. Stabilization therapy can succeed only if resistant cells are excluded and the tumor converges to a fully sensitive equilibrium. This makes the feasibility of stabilization therapy more restrictive than in models with fixed resistance levels or a consistent resistance cost [27, 28].

The fitted model also suggested that treatment can interact with initial population composition to shape resistance evolution. In the presence of alectinib, resistant cells evolved faster when they were initially rare than when they were initially frequent. One possible explanation is that treatment reduces the sensitive-cell population, creating ecological opportunity for the remaining resistant cells. In such a setting, resistant cells may experience reduced competition and greater access to resources, which can make more aggressive resistant phenotypes favorable. This interpretation is consistent with the idea that therapy changes not only population size, but also the selective environment in which resistance evolves. Because the experiment was short, the inferred changes in the resistance trait most likely reflect phenotypic plasticity, epigenetic change, or selection among pre-existing phenotypic states rather than new genetic evolution. This is important clinically, because non-genetic resistance can arise on treatment-relevant timescales and may alter response before stable genetic resistance is detected [21, 22].

A second context-dependent pattern concerned innate resistance. In the fitted model, the innate resistance parameter *k* modulates treatment-induced mortality in both sensitive and resistant cells. Higher inferred values of *k* correspond to weaker effective treatment response. We found higher inferred innate resistance in cultures where sensitive cells were initially more abundant. This result should not be interpreted as showing that sensitive cells are intrinsically more resistant. Rather, it suggests that the effective treatment response of the mixed population depends on its initial composition. One possible explanation is a shielding or dilution effect: abundant sensitive cells may absorb treatment effects or alter the ecological conditions experienced by resistant cells. More generally, this finding reinforces that fitted resistance parameters can reflect population-level eco-evolutionary effects, not only cell-intrinsic molecular mechanisms.

The parameter correlations further support the view that resistance dynamics should be interpreted at the system level. Across experimental conditions, intrinsic growth rate and natural death rate were strongly correlated, suggesting that fitted models may capture variation in cell turnover rather than growth alone. Under treatment, when sensitive cells were initially in the majority, evolutionary speed, innate resistance, and resistance benefit were positively correlated. This pattern suggests that resistance evolution, effective treatment response, and the benefit of increasing resistance may form a linked dynamical syndrome under certain ecological conditions. Such correlations are relevant for model development because they indicate that parameters may not vary independently across biological contexts. They also highlight the need for caution when interpreting a single fitted parameter as a direct biological mechanism.

Our results extend the NSCLC study of Kaznatcheev et al. [20], who showed that alectinib and fibroblasts can change the evolutionary game played by sensitive and resistant NSCLC cells. Their analysis focused on frequency-dependent interactions using a matrix-game approach. Here, we used a population-dynamic model that explicitly tracks sensitive and resistant population sizes and allows the resistant population to evolve along a continuous resistance trait. This allowed us to ask a complementary question: not only whether ecological interactions differ across conditions, but whether the fitness effect of resistance itself changes with ecological context. The answer from the fitted model is yes. The model also connects naturally to recent polymorphic cancer growth models, which show that mixed cancer populations can require more than a single homogeneous growth law to describe treatment response across *in-vitro* and *in-vivo* settings [25].

The clinical translation of these results requires care. The data are *in-vitro*, the treatment was represented by a simplified dose effect, and the fitted resistance trait is a model variable rather than a directly measured molecular phenotype. Moreover, the current analysis does not establish a clinically implementable dosing algorithm. Instead, it identifies conditions under which different evolutionary therapy principles may or may not apply. In particular, our dose simulations suggest that maximum tolerated dosing is not always the best way to maximize time to progression in the gain-of-resistance regime. When an intermediate dose keeps the initial tumor growth rate close to zero, it can delay progression longer than a higher dose in the simulated system. This should be viewed as a model-generated hypothesis, not a clinical recommendation. Its value is to show that treatment design depends on the resistance regime, and that the same dosing principle may not apply across ecological contexts.

These findings also emphasize the importance of monitoring. If the fitness effect of resistance depends on tumor composition, then estimating only total tumor burden is insufficient for evolutionary therapy. Ideally, treatment decisions would use longitudinal information about tumor burden, resistant-cell abundance, and changes in resistance phenotype. In metastatic NSCLC, this is challenging because repeated tissue biopsies are often infeasible and reliable serum biomarkers are limited. This makes model-supported integration of imaging, liquid biopsies, and other longitudinal data especially important [40–42]. Our results suggest that such monitoring should aim not only to detect resistance, but also to infer whether resistance is currently costly or beneficial in the patient-specific ecological context.

In summary, this study shows that the consequences of resistance for evolutionary therapy may depend on how resistance evolves during treatment and on the ecological composition of the tumor. In the fitted NSCLC cultures, resistance could behave as either costly or growth-promoting, depending on initial population composition. This context dependence changes the feasibility of stabilization therapy and affects which dosing principles are expected to prolong control. Evolutionary therapy for NSCLC should therefore account not only for the abundance of resistant cells, but also for how resistance changes cellular fitness and treatment response over time.

## Acknowledgments

We thank Artem Kaznatcheev for sharing the data from Kaznatcheev et al. [20], which we used to fit the models in this study.

## A Existence and stability of the equilibria in the negative resistance case

In this section, we analyze the existence and stability of equilibria in the negative resistance case, *g <* 0. The positive resistance case, *g >* 0, is analyzed in [27].

### A.1 Interior equilibrium

If an interior equilibrium 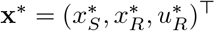 of (4) exists, then, according to 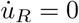, we should have:

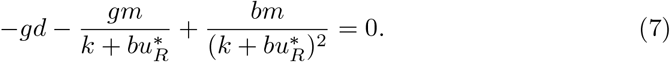

Since *g <* 0, the left-hand side of (7) is positive and therefore cannot be equal to zero. Thus, no interior equilibrium exists in this case.

### A.2 Trivial equilibrium

By a trivial equilibrium, we mean an equilibrium with *x*_*S*_ = *x*_*R*_ = 0, while 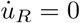, i.e.,

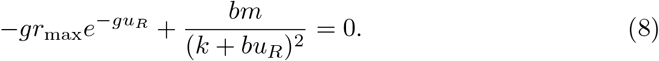

In (8), the left-hand side is positive because *g <* 0, and therefore it cannot be equal to zero. Thus, no trivial equilibrium exists in the negative resistance case.

### A.3 Fully sensitive equilibrium

At this equilibrium, *x*_*R*_ = 0. We refer to this equilibrium as 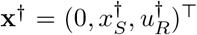. Sub-stituting *x*_*R*_ = 0 into 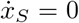 and 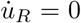 yields the following expressions for 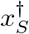 and 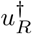:

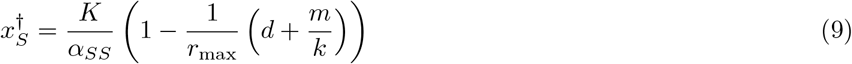

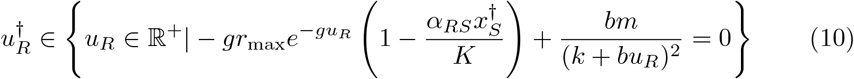

An argument similar to that used for (8) applies to (10), implying that (10) can have up to two solutions.

Since 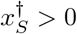, it follows that

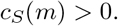

In addition, (10) implies

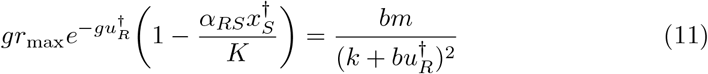

and, since *g <* 0, this leads to

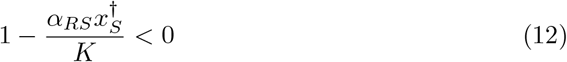

Substituting 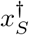 from (9) into (12), we obtain

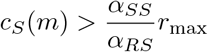

Thus, the existence of the fully sensitive equilibrium requires the inequality

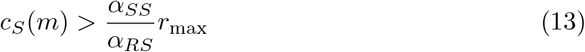

to hold.

The eigenvalues of the Jacobian matrix evaluated at **x**^*†*^ are:

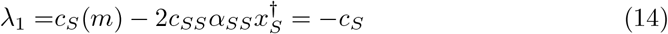

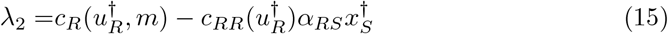

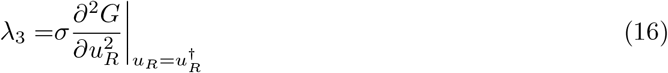

Using the equations 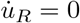 and 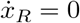, we can rewrite (16) as

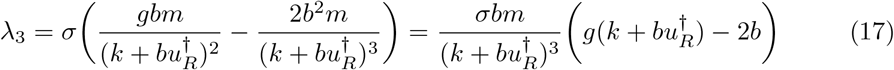

Since *g <* 0, (17) is negative, and consequently *λ*_3_ *<* 0.

Substituting (9) into (15), we can rewrite *λ*_2_ as

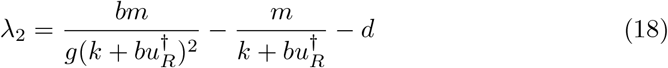

Again, since *g <* 0, *λ*_2_ is negative. As a result, all eigenvalues of the Jacobian matrix are negative. Thus, if the fully sensitive equilibrium exists, it is locally stable.

### A.4 Fully resistant equilibrium

At this equilibrium, which we denote by 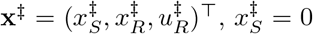, while 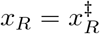 and 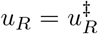, with

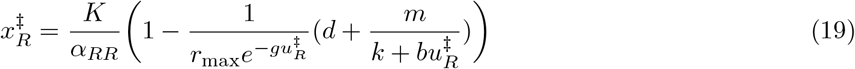

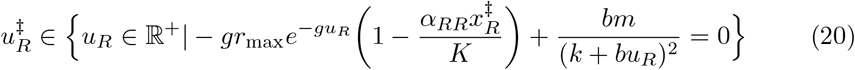

Substituting (19) into (20) yields

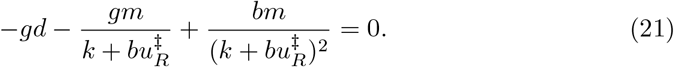

Since *g <* 0, the left-hand side of (21) is positive and therefore cannot be equal to zero. Thus, no fully resistant equilibrium exists in the negative resistance case.

## B Analyzing the existence of a Darwinian demon

In this case, according to (4), 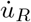 becomes negative and remains negative until it reaches zero, unless an equilibrium exists before zero is reached or a stable fully sensitive equilibrium exists, as shown in Figure 10. When *u*_*R*_ = 0, resistant cells have effectively become sensitive cells.

**Fig. 10:**
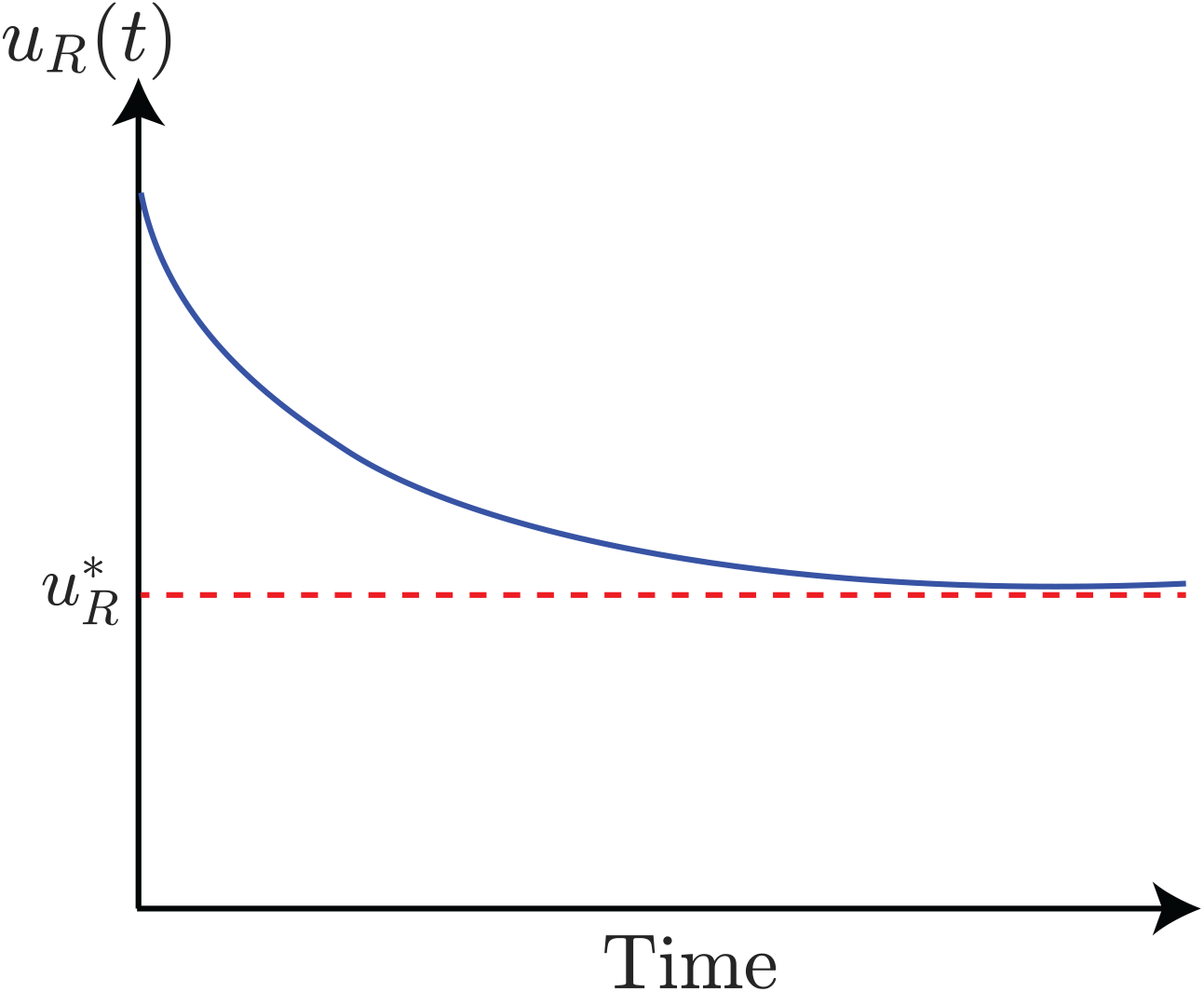
Existence of the interior equilibrium.

It was shown in the previous sections that, for an interior equilibrium to exist, we should have

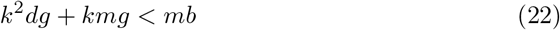

However, in this case, the left-hand side is positive and the right-hand side is negative, which is impossible. Thus, neither an interior equilibrium nor a fully resistant equilibrium can exist. For the trivial equilibrium, we showed that if *gb <* 0, this equilibrium does not exist. Moreover, for the fully sensitive equilibrium, we showed that if *gb <* 0, the population size at equilibrium exceeds the tolerable threshold for the patient. Consequently, no equilibrium exists before *u*_*R*_ reaches zero, nor does a stable fully sensitive equilibrium exist.

## C Structure of the data

The initial seeding of each well is defined using the following *I*_*S*_ matrix. The entries represent the proportion of sensitive cells at the beginning of the experiment. In addition, the presence of alectinib and fibroblasts in the experiment is indicated by the matrices *Al* and *Fb*, respectively. In both *Al* and *Fb*, 1 represents the presence and 0 represents the absence of the corresponding agent.

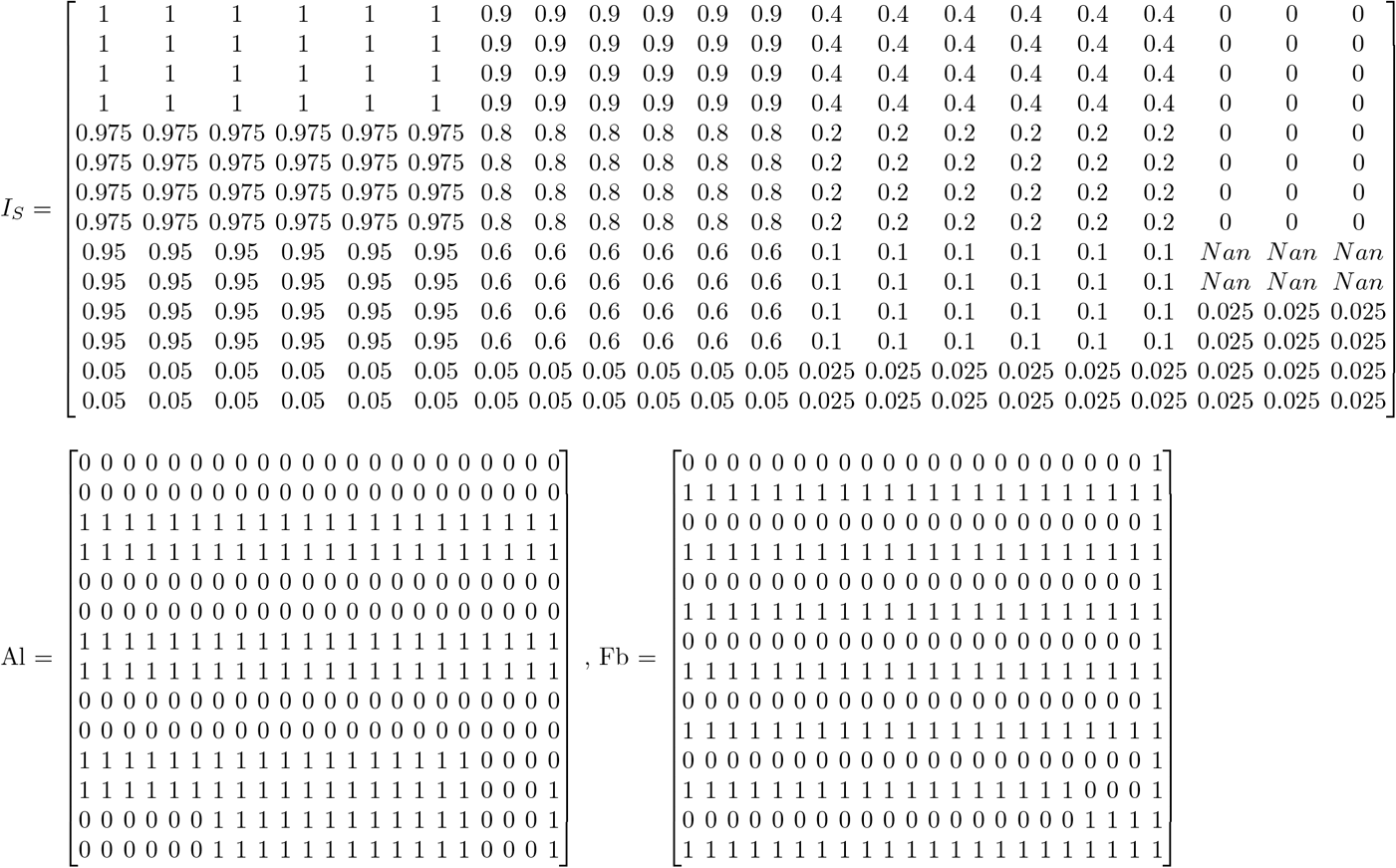

## D Fitting results

Some of the fitting results for the negative and positive resistance cases are presented in Figures 11 and 12.

**Fig. 11:**
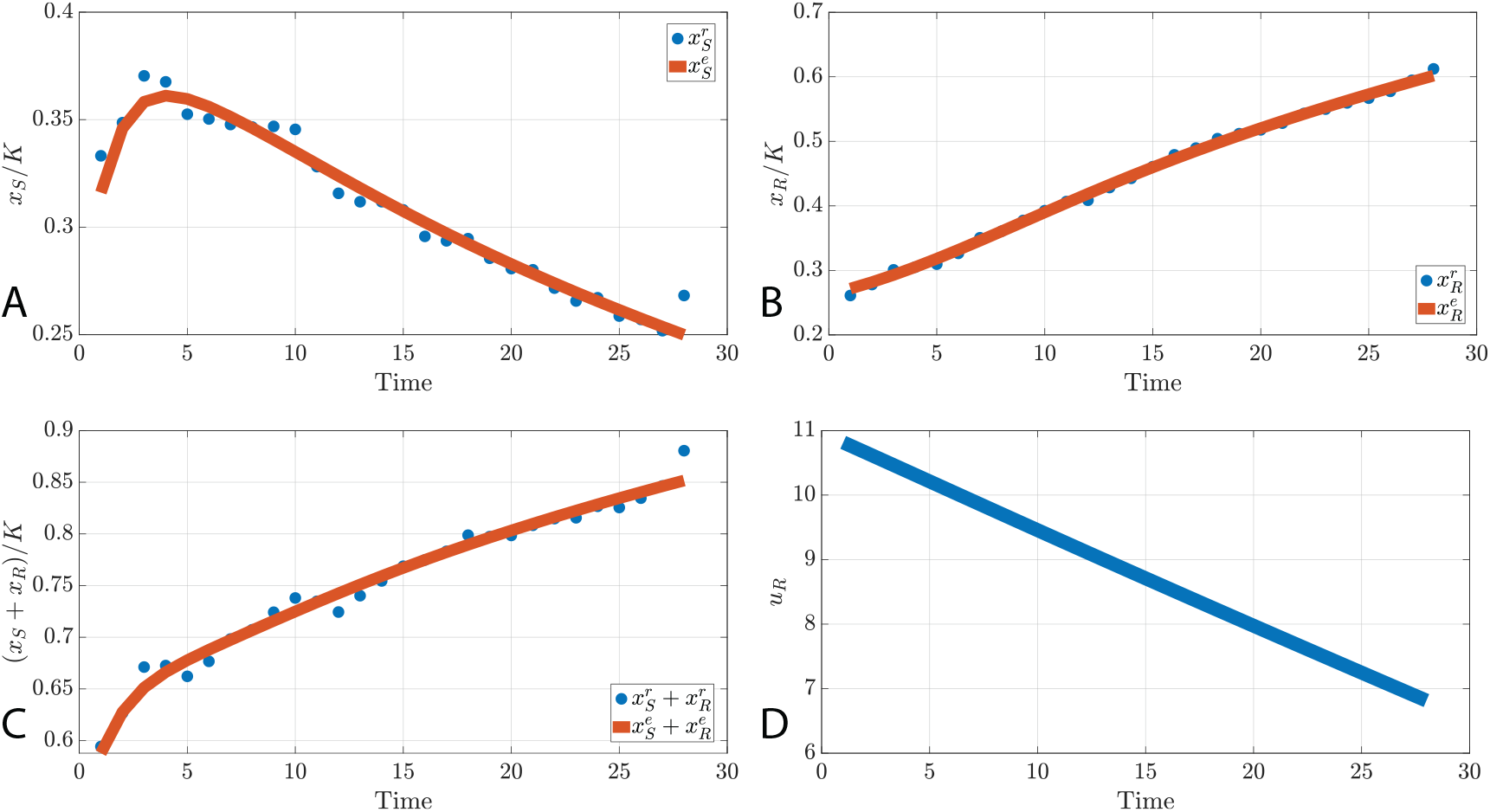
Data fitting for the well (3,18) with *g >* 0, positive resistance. A : Normalized number of sensitive cancer cells. B : Normalized number of resistant cancer cells. C : Normalized number of all cancer cells. D : Resistance rate.

**Fig. 12:**
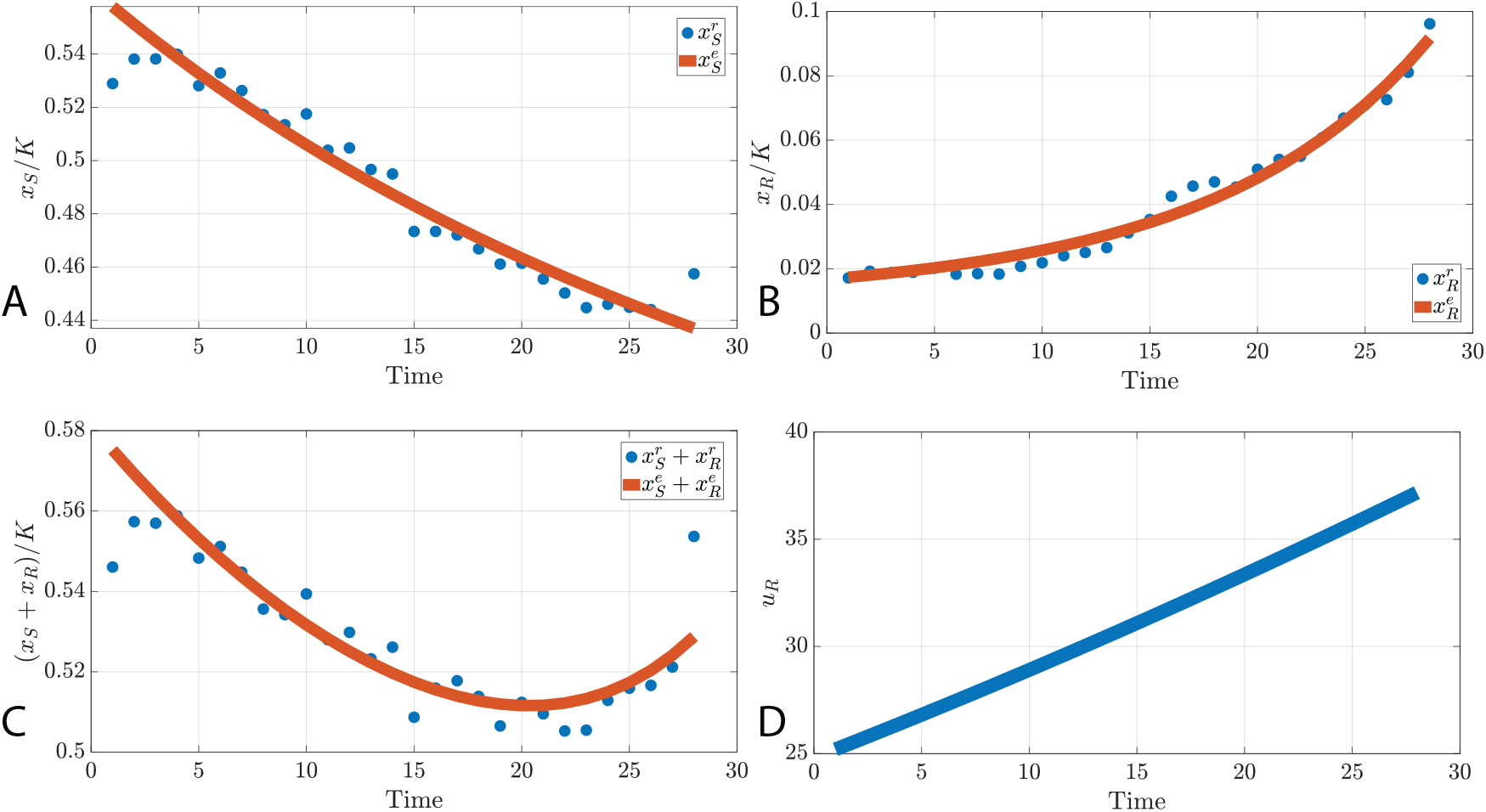
Data fitting for the well (7,6) with *g <* 0, negative resistance. A : Normalized number of sensitive cancer cells. B : Normalized number of resistant cancer cells. C : Normalized number of all cancer cells. D : Resistance rate.

## E Spearman rank correlations and scatter plots between the parameters of cancer dynamic

In this section, we show the correlation matrices represented in Figure 6 in more detail, separately and with the correlation values included. In addition, scatter plots between innate resistance, growth rate, the competition coefficient between sensitive and resistant cells, and evolutionary speed are provided in Figures 13-18.

**Fig. 13:**
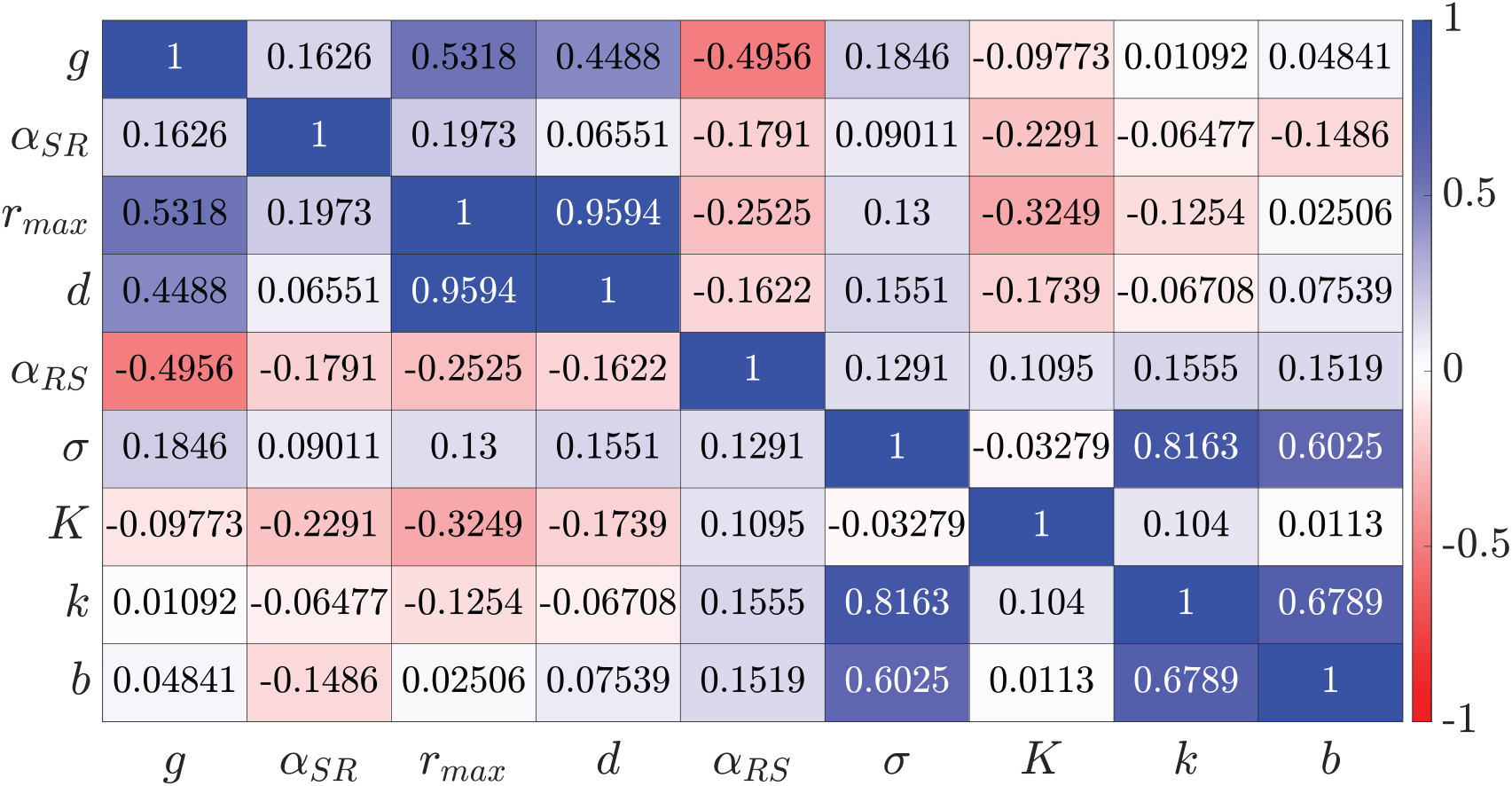
Spearman rank correlations for the case with Alectinib and starting with more sensitive cells.

**Fig. 14:**
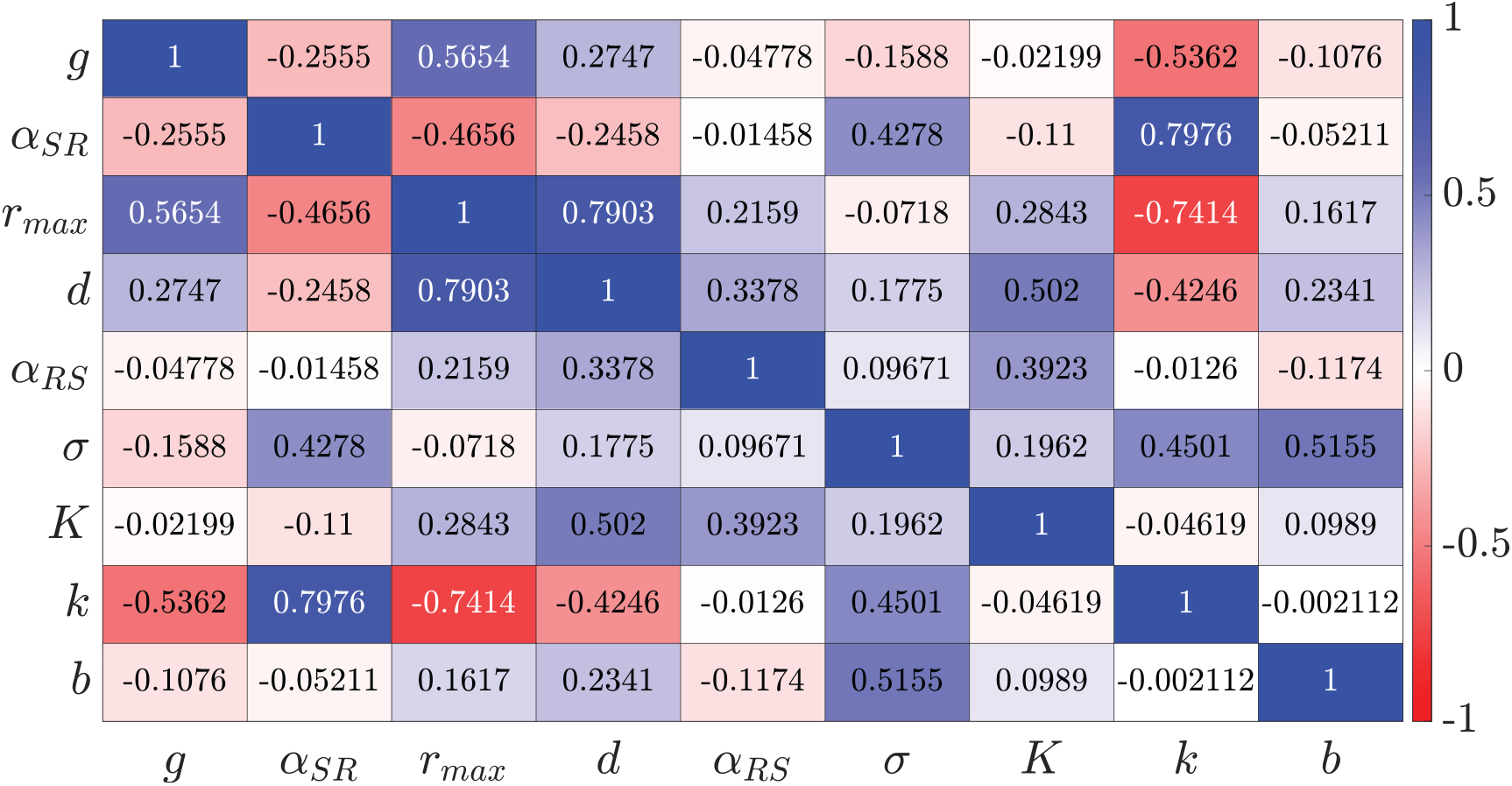
Spearman rank correlations for the case with Alectinib and starting with more resistant cells.

**Fig. 15:**
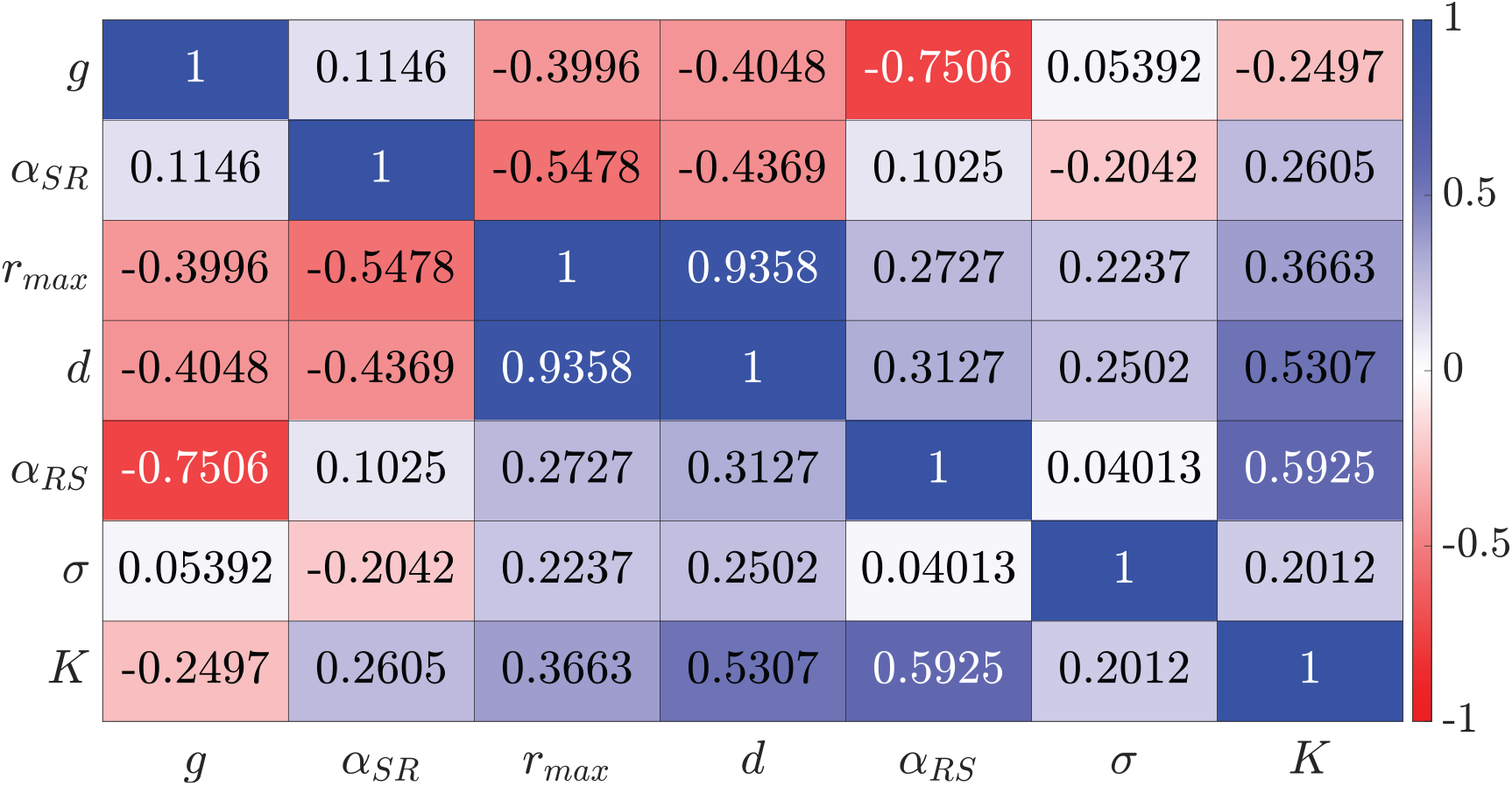
Spearman rank correlations for the case without Alectinib and starting with more sensitive cells.

**Fig. 16:**
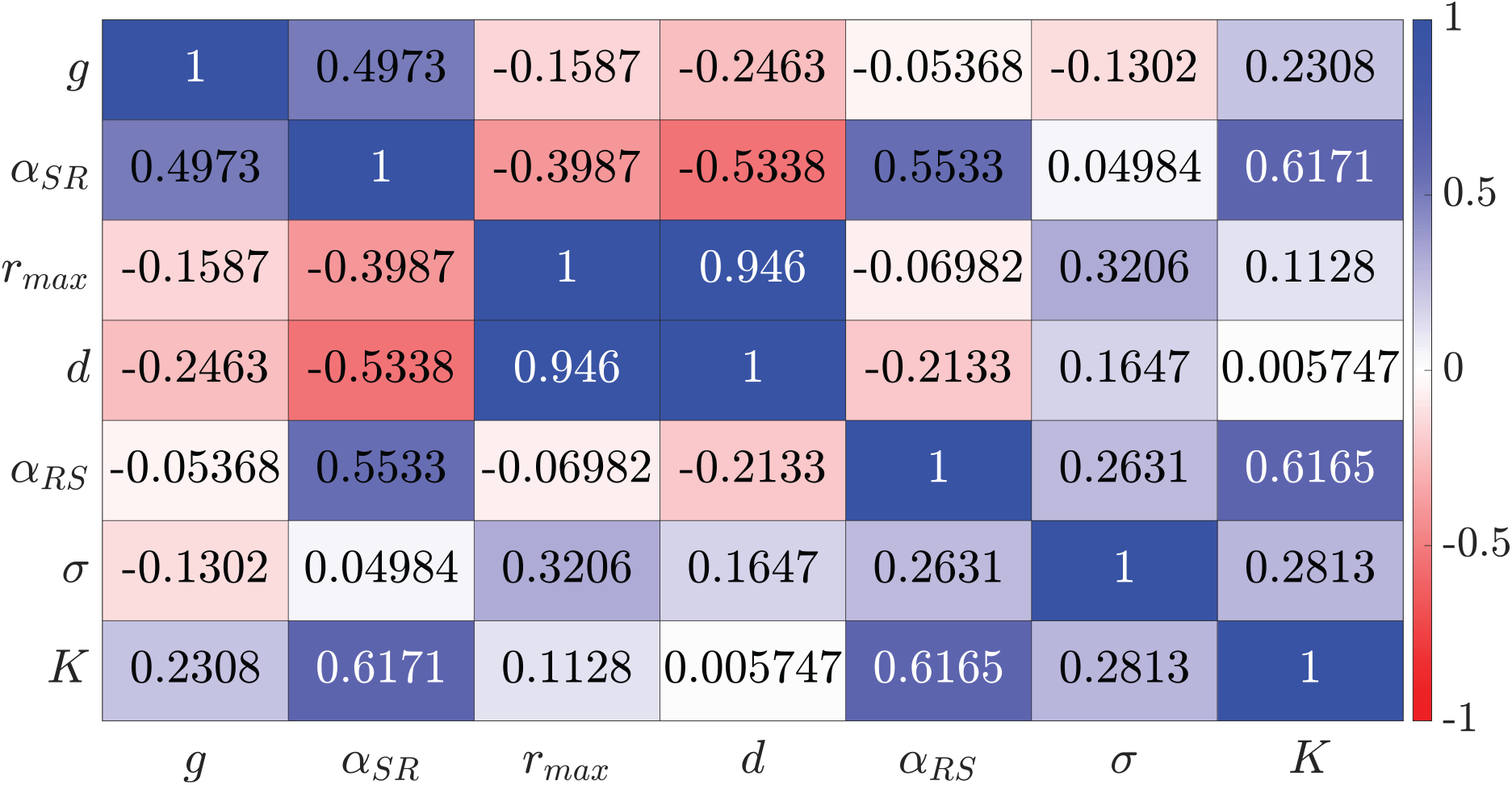
Spearman rank correlations for the case without Alectinib and starting with more resistant cells.

**Fig. 17:**
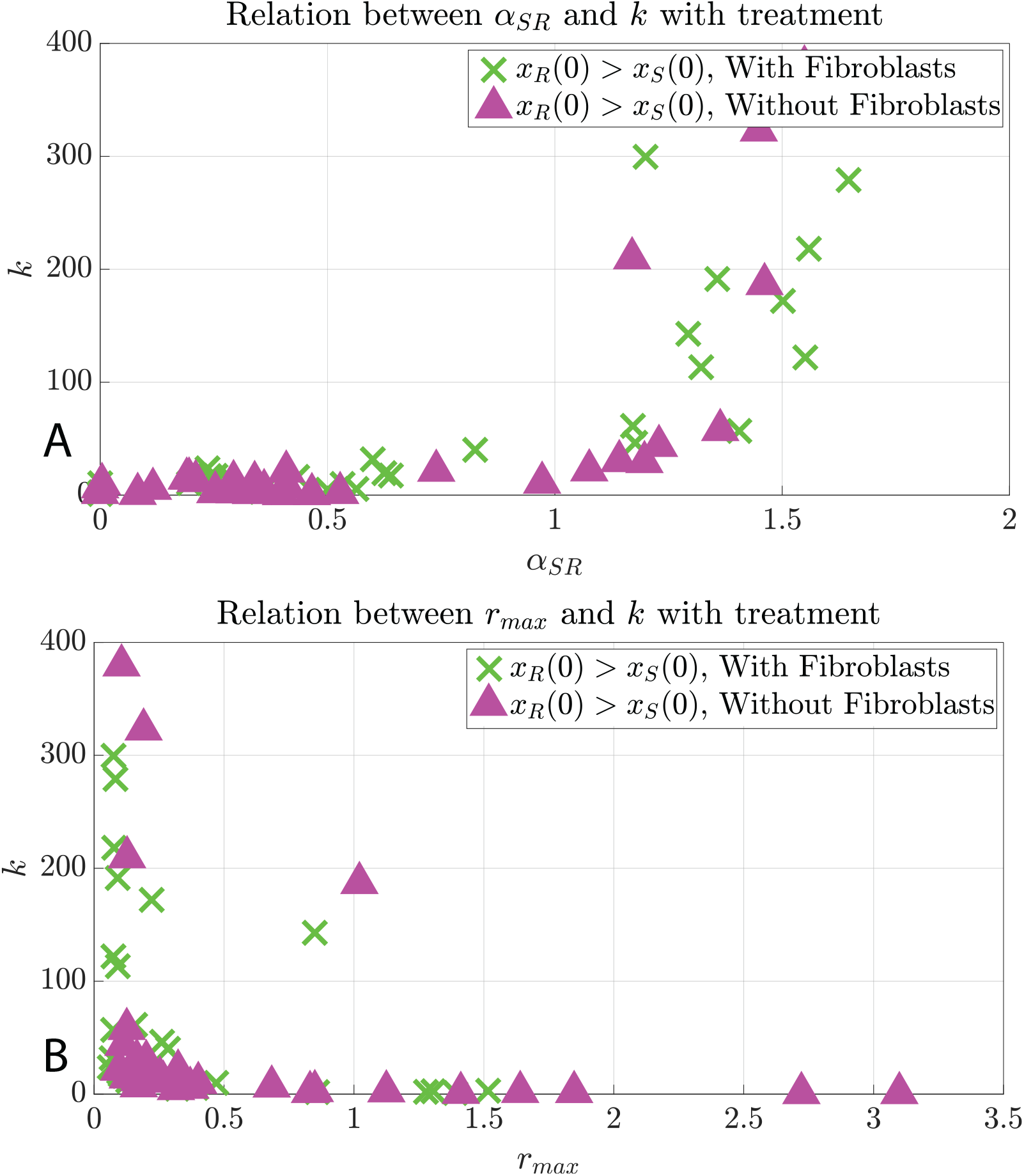
Scatter plots between innate resistance, growth rate, and one of the competition coefficients in the presence of Alectinib.

**Fig. 18:**
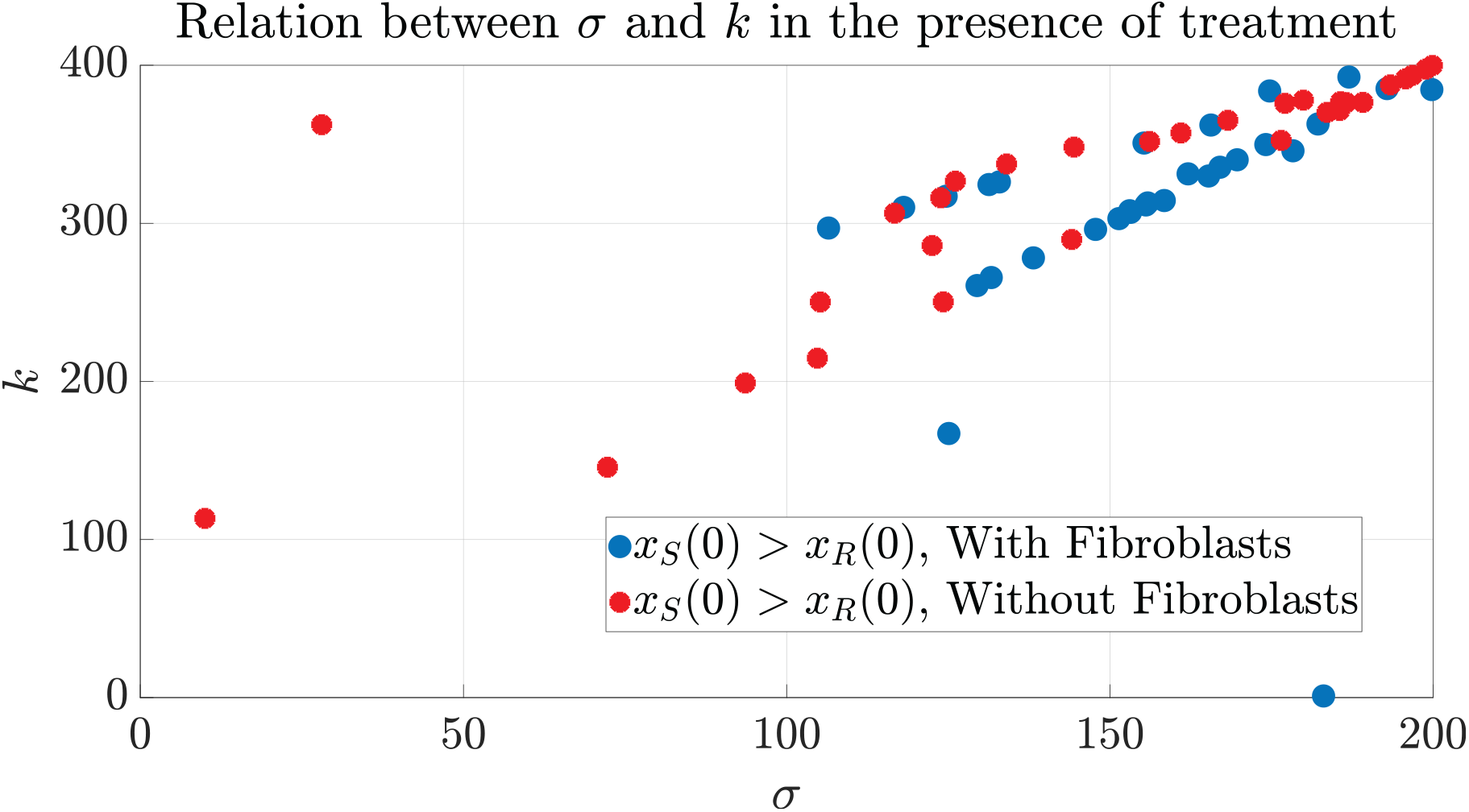
Scatter plot between innate resistance and evolutionary speed. These two parameters are monotonically correlated.

